# Semaphorin 3A induces cytoskeletal paralysis in tumor-specific CD8+ T cells

**DOI:** 10.1101/849083

**Authors:** Mike B Barnkob, Yale S Michaels, Violaine André, Philip S Macklin, Uzi Gileadi, Salvatore Valvo, Margarida Rei, Corinna Kulicke, Ji-Li Chen, Vitul Jain, Victoria Woodcock, Huw Colin-York, Andreas V Hadjinicolaou, Youxin Kong, Viveka Mayya, Joshua A Bull, Pramila Rijal, Christopher W Pugh, Alain R Townsend, Lars R Olsen, Marco Fritzsche, Tudor A Fulga, Michael L Dustin, E Yvonne Jones, Vincenzo Cerundolo

**Affiliations:** MRC Human Immunology Unit, MRC Weatherall Institute of Molecular Medicine, University of Oxford, Headley Way, Oxford OX3 9DS, UK; MRC Weatherall Institute of Molecular Medicine, University of Oxford, Headley Way, Oxford OX3 9DS, UK; Nuffield Department of Medicine, University of Oxford, Nuffield Department of Medicine Research Building, Roosevelt Drive, Oxford OX3 7FZ, UK; Kennedy Institute of Rheumatology, University of Oxford, Roosevelt Dr, Oxford OX3 7FY, UK; Division of Structural Biology, Wellcome Centre for Human Genetics, University of Oxford, Roosevelt Drive, Oxford, OX3 7BN, UK; Wolfson Centre for Mathematical Biology, Mathematical Institute, University of Oxford, Radcliffe Observatory Quarter, Woodstock Road, Oxford, OX2 6GG, UK; Department of Health Technology, Technical University of Denmark, Ørsteds Plads, Building 345C, 2800 Kgs. Lyngby, Denmark

**Keywords:** cancer, immune system, T cells, semaphorin, neuropilin-1, immunotherapy, cytoskeleton

## Abstract

Semaphorin-3A (Sema3A) regulates tumor angiogenesis, but its role in modulating anti-tumor immunity is unclear. We demonstrate that Sema3A secreted within the tumor microenvironment (TME) suppresses tumor-specific CD8+ T cell function via Neuropilin-1 (NRP1), a receptor that is upregulated upon activation with T cells’ cognate antigen. Sema3A inhibits T cell migration, assembly of the immunological synapse, and tumor killing. It achieves these functional effects through hyper-activating the acto-myosin system in T cells leading to cellular paralysis. Finally, using a clear cell renal cell carcinoma patient cohort, we demonstrate that human tumor-specific CD8+ T cells express NRP1 and are trapped in Sema3A rich regions of tumors. Our study establishes Sema3A as a potent inhibitor of anti-tumor immunity.

## INTRODUCTION

Cytotoxic CD8+ T cells are often restricted to certain areas within tumors or completely excluded from the tumor microenvironment (TME) (*1*). We hypothesized that cell guidance cues involved in developmental processes may also play a role in T cell restriction in the tumor microenvironment. The secreted protein Sema3A is known to guide both endothelial cells and neurons during embryogenesis through the cell-surface receptor family Plexin-A (*2, 3*). Sema3A binding to Plexin-A requires the co-receptor NRP1 (*4, 5*). In axonal growth cones, Sema3A signaling leads to profound changes in filamentous actin (F-actin) cytoskeletal organization (*6*), an effect that is thought to be dependent on myosin-IIA activity (*7*). Sema3A can also be produced by cancer cells (*8*) and recent evidence indicates that NRP1, like PD-1, is upregulated on dysfunctional tumor-specific CD8+ T cells and can modulate their anti-tumor response (*9–11*). However, there is no consensus on whether the Sema3A-NRP1 axis is immunosuppressive (*8, 12*) or supportive of CD8+ T cells’ response to tumors (*13*). Furthermore, due to Sema3A’s anti-angiogenic effects (*14*), several groups have proposed utilizing Sema3A to inhibit tumor growth (*13, 15*). It is therefore critical to examine the role of Sema3A in anti-tumor immunity more closely.

## RESULTS

### Tumor-specific CD8+ T cells upregulate NRP1 and Plexin-A1

To establish whether Sema3A can affect CD8+ T cells, we first examined NRP1 expression of its cognate receptor, NRP1 on naive and stimulated T cells. NRP1 was upregulated on human NY-ESO-1-specific HLA-A2 restricted CD8+ T cells, as well as on murine OT-I CD8+ T cells (OT-I T cells), upon stimulation with their cognate peptides, NY-ESO-1_157-165_ and Ovalbumin_257-264_ (Ova), respectively (**Figure 1A-B**). Analysis of transcriptional data from the Immunological Genome Project Consortium (*16*) of naive and effector CD8+ T cells corroborated these findings (**Supplementary Figure 1A**). We examined whole OT-I T-cell protein lysate and found that two NRP1 isoforms exist in murine T cells, with the larger NRP1 protein being the dominant form following T cell activation (**Supplementary Figure 1B**). To examine NRP1 regulation in CD8+ T cells, we utilized antigenic Ova peptides with varying affinities for the OT-I TCR (*17*), namely SIINFEKL (N4), SIIQFEKL (Q4) and SIITFEKL (T4) and found that NRP1 expression was correlated with both peptide concentration and affinity of TCR engagement (**Figure 1C**).

**Figure 1.**
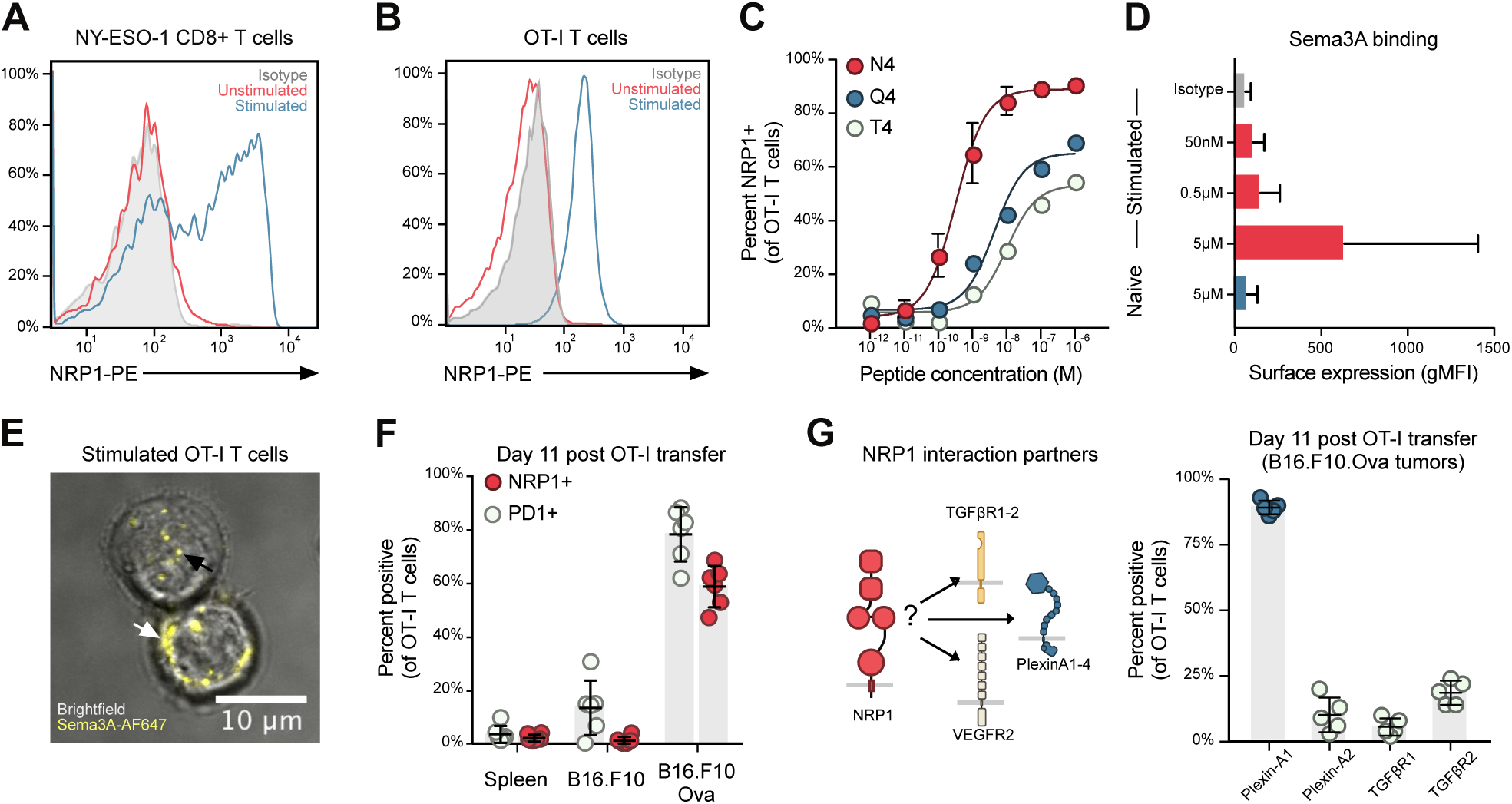
Tumor-specific CD8+ T cells up-regulate NRP1 and Plexin-A1 allowing for Sema3A binding. **A-B**. Representative histogram of flow cytometric analysis of surface NRP1 expression on human NY-ESO-1-specific HLA-A2 restricted CD8+ T cells and murine OT-I CD8+ T cells following 48 hours stimulation with cognate peptides. Cells are gated on CD45, CD8 and TCRβ. Experiment repeated three times. **C.** Analysis of NRP1 up-regulation using peptides with varying TCR affinities. Cells are gated on CD45.1, CD8 and TCRβ. Cells from 3 mice per group, experiment was performed once. Data indicate mean ± SD. **D.** Quantification of surface binding of Sema3A_S-P_ on naïve and 48 hour stimulated OT-I T cells. Cells are gated on CD8 and CD3. Experiment was repeated three times. Data indicate mean ± SD of representative experiment. **E.** Confocal imaging of 48 hour stimulated OT-I T cells stained with AF647-labelled Sema3A_S-P_ shows that the protein can bind to the cell membrane (white arrow) and within the cell (black arrow). **F.** Flow cytometric analysis of PD-1 and NRP1 expression on OT-I T cells 11 days after adoptive transfer in spleen, non-antigen expressing tumor (B16.F10) and antigen-expressing tumor (B16.F10.Ova) (n=6). Data representative of two independent experiments and indicate mean ± SD of six mice per group. **G.** Schematic of NRP1 interactions partners (left). Flow cytometric analysis of expression of selected NRP1 interactions partners on OT-I T cells 11 days after adoptive transfer (n=5) (right). Experiment was performed once. Data indicate mean ± SD. Abbreviations: gMFI, geometric mean fluorescence intensity. N4, SIINFEKL. Q4, SIIQFEKL. T4, SIITFEKL.

NRP1 is a co-receptor for a number of cell-surface receptors, including TGFβ receptors 1 and 2 (TGFβR1-2) (*18*), VEGF receptor 2 (VEGFR2) (*19*) and Plexin-A1, -A2, -A3 and -A4 receptors (*20*), and its function is highly dependent on the availability of these receptors for downstream signaling. We therefore screened OT-I T cells for expression of NRP1 partner receptors. Stimulated, but not naive, OT-I T cells expressed Plexin-A1 but little to no Plexin-A2, TGFβR1, TGFβR2 or VEGFR2 (**Supplementary Figure 1C-E**). Plexin-A4 was expressed at low levels on both unstimulated and stimulated cells. Analysis of Plexin-A3 expression was not included because antibodies specific to Plexin-A3 could not be found. Having identified NRP1 and Plexin-A1 receptors on stimulated, but not naive T cells, we expected Sema3A ligation to the former (*5*). Indeed, flow cytometric analysis confirmed that only stimulated OT-I T cells could bind recombinant murine Sema3A_S-P_ (**Figure 1D**). Confocal imaging further indicated that Sema3A was internalized upon binding to T cells (**Figure 1E**).

We next explored whether NRP1 and Plexin-A1 expression would be retained by CD8+ T cells during infiltration in the TME. We adoptively transferred congenically marked and activated OT-I T cells into syngeneic C57BL/6 mice bearing either B16.F10 or OVA expressing B16.F10 cells (B16.F10.Ova) in opposing flanks. While few NRP1 expressing OT-I T cells infiltrating B16.F10 control tumors were NRP1 positive, the majority of OT-I T cells residing within B16.F10.Ova tumors expressed NRP1 (**Figure 1F**) and Plexin-A1 (**Figure 1G, right**) up to eleven days after adoptive transfer. Of note, endogenous CD4+CD25+FoxP3+ T cells found within the tumor expressed both NRP1 and Plexin-A1 as well as TGFβR1-2 (**Supplementary Figure 1E**), indicating that this subset of T cells might be modulated differently from CD8+ T cells. Collectively, these data show that NRP1 and Plexin-A1 receptors are upregulated on CD8+ T cells in a TCR-dependent manner, that they are expressed on tumor-specific OT-I T cells, and that recombinant Sema3A can bind directly to activated CD8+ T cells.

### Sema3A negatively regulates CD8+ T cell adhesion, motility and migration through NRP1

Sema3A is known to restrict neuronal migration (*4*), but can have opposing effects on immune cell motility. While both thymocyte (*21*) and macrophage (*22*) migration can be inhibited, Sema3A has also been shown to increase dendritic cell (DC) migration (*23*). We therefore undertook a number of *in vitro* experiments designed to dissect the effect of Sema3A on CD8+ T cell adhesion and motility. We first utilized interference reflection microscopy (IRM) to assess T cell contact and adhesion (*24*). This was done on plates coated with ICAM-1 and the chemokine ligand C-X-C motif chemokine ligand 12 (CXCL12, SDF-1a) in order to emulate the environment found on endothelial cells and extracellular matrix within the TME (*25*). When Sema3A_S-P_ was coated on plates, T cell adhesion was significantly weakened (**Figure 2A**), an effect that was present from initial attachment until at least 10 minutes later (**Figure 2B**). In addition T cells displayed a reduced polarized morphology (**Figure 2C, Supplementary Figure 2A**). T cell motility was also affected, as both distance and velocity were reduced when Sema3A_S-P_ was present, an effect that could be reverted by pre-treating T cells with a blocking anti-NRP1-antibody (**Figure 2D-E**). Extravasation into tumors requires T cells to first adhere to endothelial cells and then transmigrate into the underlying parenchyma. To model this, we performed two experiments. First, we perfused T cells across surfaces with ICAM-1 and CXCL12 with or without Sema3A_S-P_, and found that under a range of external flow rates, Sema3A decreased the number of cells able to display rolling or tight adhesion (**Figure 2F-G**). At flow rates of 80 μm/sec, many T cells had a migration path similar to laminar flow indicating little ability to adhere (**Figure 2F, lower right figure**). Secondly, using a transwell assay, we found that Sema3A strongly inhibited transmigration (**Figure 2H**). We wondered if these effects were mediated through changed expression levels of integrins or selectins involved in adhesion and extravasation. However, flow cytometric analysis did not reveal any down-regulation of CD11a (part of LFA-1), CD49d or CD162 (**Supplementary Figure 2B**), suggesting that Sema3A signaling does not affect expression of these archetypal adhesion receptors on CD8+ T cells. These data illustrate that Sema3A strongly inhibits activated CD8+ T cell adhesion and motility, an effect that can be modulated using anti-NRP1-blocking antibodies.

**Figure 2.**
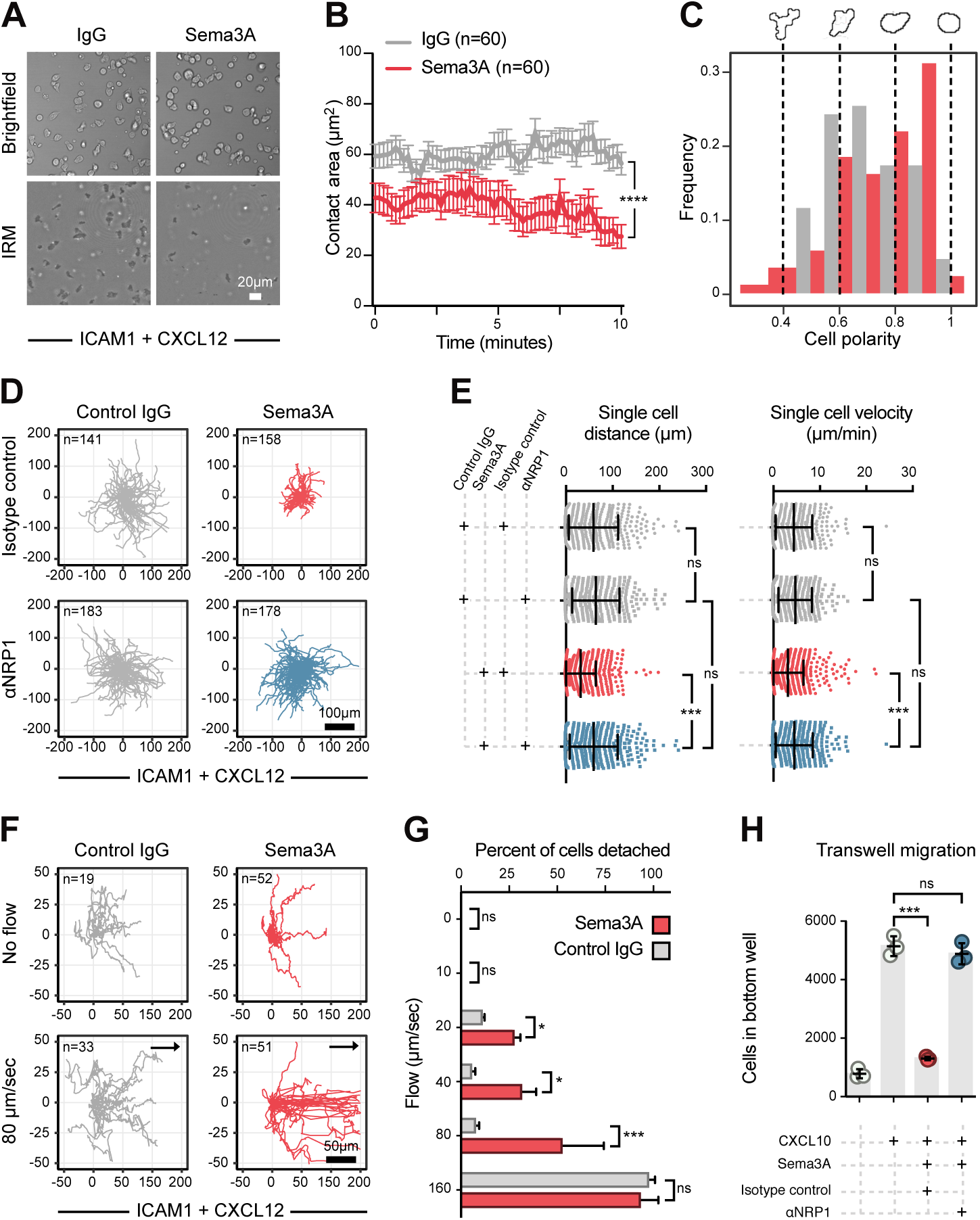
Sema3A negatively regulates CD8+ T cell adhesion, motility and migration through NRP1. **A.** Representative brightfield and IRM images of 48 hour stimulated OT-I T cells adhering to ICAM-1 and CXCL12 coated plates with either Sema3A_S-P_ or IgG immobilized. **B.** Quantification of contact area per single cell using live-cell microscopy for 10 minutes after OT-I T cells were added to plate (n=60 cells). Data representative of three independent experiments and indicate mean ± SEM. **** = P < 0.0001 by Student’s t-test. **C.** Relative frequency of cell polarity from brightfield images. A polarity of 1 indicates a shape of a perfect circle, 0 a rectangular shape. Representative images of OT-I T cells illustrated above graph. **D.** Representative spider plots showing the migration paths of individual T cells pre-treated with either a NRP1 blocking antibody or isotype control antibody on similar plates as in (A). **E.** Graph of single cell distance (left) and single cell velocity (right) in same experiment as (D). (n=314-744 cells per group). Data combined from five independent experiments indicate mean ± SD. *** = P < 0.001, ns = not significant by Kruskal-Wallis test. **F.** Representative spider plots showing the migration path of individual OT-I T cells on similar plates as in (A), with flow rates at 0 or 80 μm/sec. Arrows indicate flow direction. **G.** Quantification of percent of OT-I cells that detach in same experiment as (F) (n=20-73 cells per condition). Data representative of two independent experiments and indicate mean ± SEM. * = P < 0.05, *** = P < 0.001, ns = not significant by two-way ANOVA. **H.** Representative graph of number of stimulated OT-I T cells able to transmigrate through 3 μm Boyden chamber with CXCL12 in bottom chamber, with or without Sema3A_S-P_ in top-chamber. OT-I T cells were pre-treated with either a blocking NRP1 antibody or isotype control antibody. Data representative of two independent experiments and indicate mean ± SD. *** = P < 0.001, ns = not significant, by two-way ANOVA. Abbreviations: IRM, interference reflection microscopy. Sec, second.

### Sema3A negatively regulates CD8+ T cells’ immunological synapse formation and cell-cell contact

Given the strong effects of Sema3A on CD8+ T cell adhesion and motility, we investigated whether Sema3A also affects the formation of the immunological synapse (IS). We first tested the ability of CD8+ T cells to form close contacts with an activating surface displaying immobilized ICAM-1 and anti-CD3 antibodies. To mimic an environment in which Sema3A had been secreted, T cells were added and allowed to settle in medium containing either Sema3A_S-P_ or control IgG, while the size and spreading speed of contact areas was measured using time-lapse IRM. T cells added to Sema3A-rich medium formed fewer and smaller contact zones (**Figure 3A, left, Movie S1-2**). We noticed that cells in Sema3A-rich medium did not spread as much and were slower to adhere (**Figure 3A, right**). Indeed, when analyzing contact zones over time, many cells in Sema3A-rich medium could not form large contact areas (**Figure 3B, top**) and spread at a reduced velocity (**Figure 3B, bottom**). These results were reminiscent of the effects seen when T cells were added to plates coated with ICAM-1, CXCL12 and Sema3A_S-P_ (**Figure 2A**) and indicated that T cells’ ability to form IS could be compromised as well.

**Figure 3.**
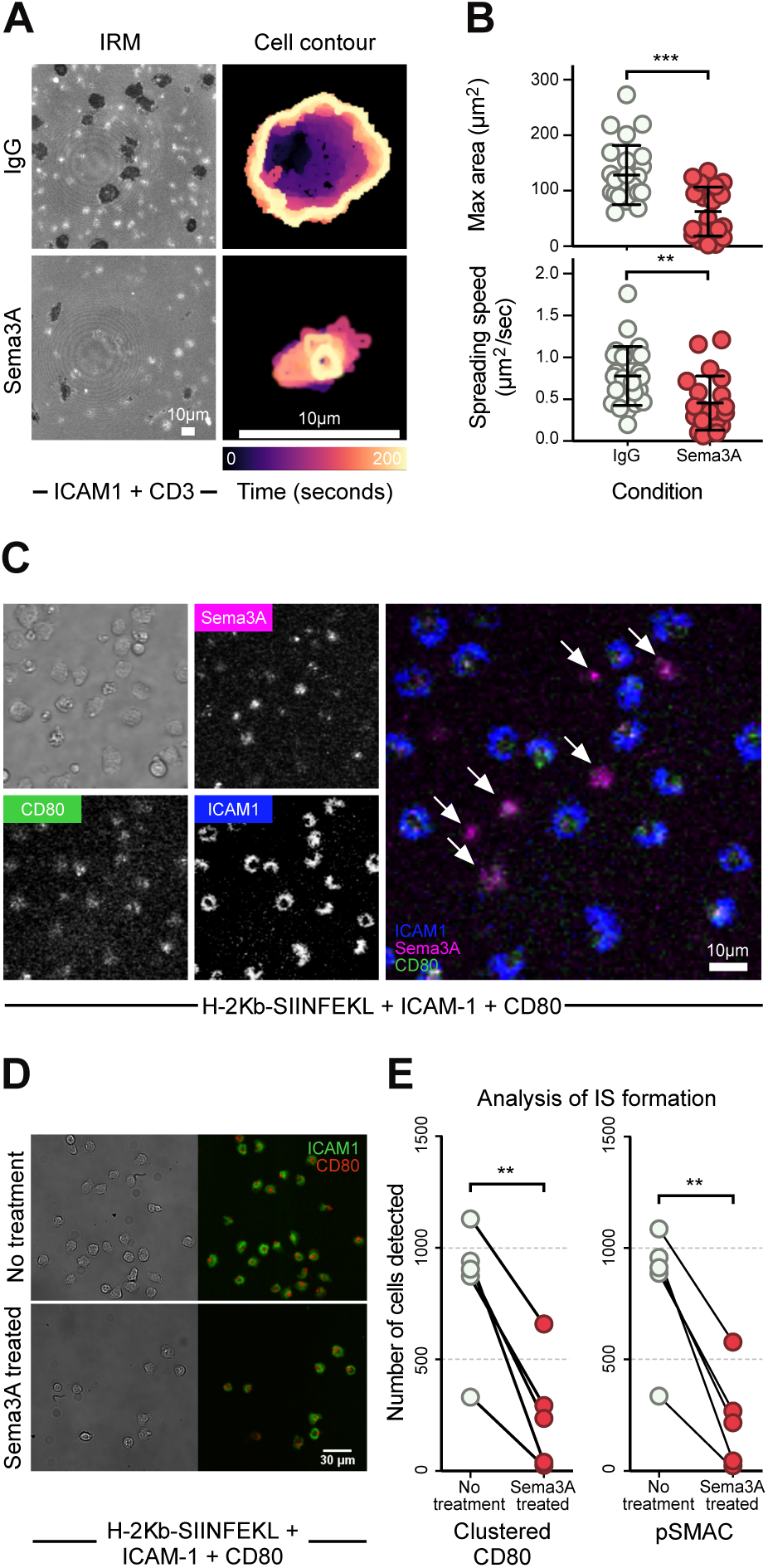
Sema3A negatively regulates CD8+ T cells’ immunological synapse formation. **A.** Live-cell imaging visualizing surface interface using IRM of stimulated CD8+ T cells dropped on an activating surface with immobilized ICAM-1 and CD3 and Sema3A_S-P_ or IgG present in medium (left). Cell contour of representative cells from either condition (right). Color of contour indicates time from 0 to 200 sec as denoted on scalebar. **B.** Quantification of maximum size of cell contact area (top) and spreading speed from initial contact to maximum contact area (bottom) (n=25 cells per group) in same experiment as (A). Data combined from three independent experiments and indicate mean ± SD. ** = P < 0.01, *** = P < 0.001, by Mann-Whitney test. **C.** Live-cell imaging of activated T cells pre-treated with Sema3A_S-P-I_-AF647 and allowed to form synapses on supported lipid bilayers with ICAM-1, CD80 and H-2Kb-SIINFEKL. Arrows in merged image indicate cells that have bound Sema3A and do not form immunological synapses. **D.** Representative image from high-throughput analysis of immunological synapses on supported lipid bilayers as in (C) with OT-I T cells pre-treated with Sema3A or not. **E.** Quantification of immunological synapses with or without Sema3A_S-P-I_ pre-treated OT-I T cells. Data from six independent experiments (n=90-1100 cells per mouse per group). ** = P < 0.01, by paired t-test. Abbreviations: IRM, interference reflection microscopy. Sec, seconds.

To more closely examine the effects of Sema3A on IS formation, we utilized supported lipid bilayers containing ICAM-1, CD80 and H-2 K^b^-Ova pMHC monomers. Stimulated OT-I T cells were pretreated with fluorescently-labelled Sema3A_S-P-I_, washed to ensure that residual protein did not interfere with the bilayer, and IS formation visualized using time-lapse total internal reflection fluorescence (TIRF) microscopy. T cells with none to little Sema3A-binding were seen to form classical IS containing a CD80-clustered central supramolecular activation cluster (cSMAC) and an outer ICAM-1-rich peripheral supramolecular activation cluster (pSMAC), while T cells that had strongly bound Sema3A_S-P-I_ were unable to spread and appeared incapable of engaging with CD80 and ICAM-1 on the bilayer (**Figure 3C, Movie S3**). To quantify the extent of this defect, we turned to a recently developed high-throughput method to quantify relevant IS parameters (*26*), where T cells are first fixed on the bilayer, then washed to remove non-adherent cells. Nearly two-thirds of stimulated T cells were either washed away, could not cluster CD80, or form pSMACs when pre-treated with Sema3A_S-P-I_ compared to untreated T cells (**Figure 3D-E**). Diminished IS formation in the presence of Sema3A mirrored a scenario where OT-I T cells were presented to an irrelevant pMHC-ligand, H-2 K^d^-gp33, on the bilayer (**Supplementary Figure 3A**). Among the Sema3A-treated T cells that formed IS, there was a discernible reduction in CD80 accumulation and in the radial symmetry of the synapse (**Supplementary Figure 3B-D**), indicating that CD8+ T cells can be rendered non-responsive to their cognate antigen through Sema3A signaling. We confirmed these findings by examining T cell binding to live cancer cells. Stimulated OT-I T cells and B16.F10.Ova cells were co-incubated in the presence of control IgG, Sema3A_S-P_ or a mutated Sema3A protein, in which the NRP1 interaction site on Sema3A has been mutated to substantially reduce the binding affinity (*5*), followed by enumeration of OT-I T cell:B16.F10.Ova cell-cell conjugates. We noticed a 50% reduction in number of OT-I cells capable of binding to antigen-expressing cancer cells in the presence of Sema3A, but not with the control or mutant Sema3A (**Supplementary Figure 3E**). These results thus demonstrate that Sema3A signaling leads to profound effects on a majority of CD8+ T cells’ abilities to adhere to target cells and form an IS.

### Sema3A affects T cell actin dynamics through actomyosin II activity

Class 3 semaphorins have been shown to have various effects on the cytoskeleton in hematopoietic cells, including thymocytes (*27*), dendritic cells (*23*) and T cells (*12*), however the precise nature of these effects in CD8+ T cells is not well characterized. Since cytoskeletal F-actin remodeling is necessary for T cell binding to target cells (*28*) as well as lamellopodium (*24*) and IS formation (*29, 30*), we examined F-actin content and dynamics in T cells during Sema3A_S-P_ exposure. We first treated stimulated OT-I T cells with Sema3A_S-P_ at varying durations and examined F-actin content using flow cytometry. Surprisingly, no actin depolymerization was observed up to 30 minutes after Sema3A_S-P_ treatment (**Figure 4A**). To better visualize F-actin dynamics before and after Sema3A_S-P_ treatment, we crossed LifeAct-eGFR (*31*) mice with OT-I mice to generate LifeAct-OT-I T cells. Mice developed normally and generated Ova-specific T cells with GFP-labelled F-actin. Stimulated T cells formed an active lamellopodium that undulated across an activating surface containing CD3 and ICAM-1, allowing for close inspection of F-actin dynamics using time-lapse confocal microscopy. When Sema3A_S-P_ was added to cells during this undulating phase, T cell morphology changed and took a more irregular and roughened appearance (**Figure 4B**). During this phase, F-actin content at the surface interface did not change, but lamellopodia formation stopped and F-actin became non-dynamic and immobile (**Figure 4C, 4F, Movie S4**). We therefore analyzed F-actin velocity along the cell edge using kymographs (**Figure 4D**). Sema3a_S-P_ profoundly inhibited F-actin dynamics (mean velocity was 1.34 µm/min after treatment versus 3.8 µm/min before) (**Figure 4E**). Next, we treated T cells with mutant Sema3A and found no difference in F-actin dynamics after treatment (**Figure 4E**), confirming that the effect of Sema3A on F-actin in the lamellopodia is NRP1-dependent. We assessed if this stark effect was due to localized F-actin depolymerization at the interface. Consistent with our flow cytometric analysis of global F-actin abundance (**Figure 4A**) however, the fluorescence intensity of LifeAct at the interface did not change, although the F-actin network contracted, and the cell width shrank substantially following treatment with Sema3A_S-P_ (**Figure 4F-G**). Because these effects on the actin cytoskeleton suggested that F-actin turnover dynamics could be affected, we tested wheather Jasplakinolide treatment would phenocopy the effects of Sema3A_S-Pl_. However, this instead led to constant shrinking of the cells’ F-actin network, not the immobilizing effects Sema3A produced (**Figure 4G**).

**Figure 4.**
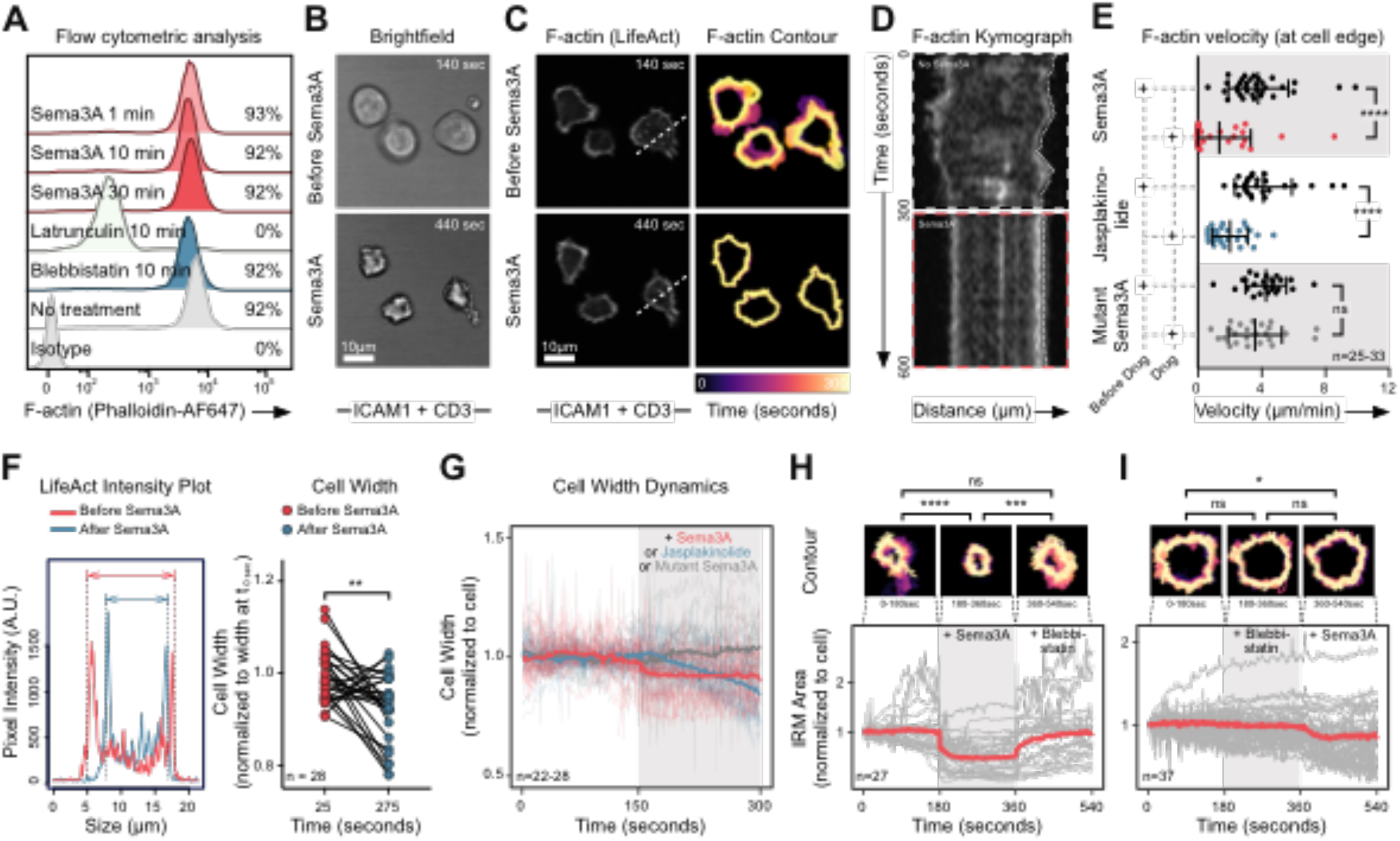
Sema3A affects T cell actin dynamics through actomyosin II activity. **A.** Representative flow cytometric analysis of F-actin content with no or varying exposure to Sema3A_S-P_ treatment in 48 hour stimulated OT-I T cells as measured by Phalloidin-staining. Percentage indicate positive cells in each condition. Data representative of two independent experiments. **B.** Representative brightfield images of 48 hour stimulated LifeAct OT-I T cells adhering to ICAM-1 and CD3 coated plates before and after Sema3A_S-P_ added to medium. **C.** Representative confocal images of LifeAct in OT-I T cells (left) and their contour plots (right) from same experiment as in (B). Image taken at cell-surface interface. Dashed white line indicate area used for (D). Color of contour indicates time from 0 to 300 sec as denoted on scalebar. **D.** Kymograph before (top) and after (bottom) Sema3A_S-P_ added to medium on area indicated with white dashed line in (C). Dotted line along edge of cell denoted example of data used for calculating data in (E). **E.** Quantification of F-actin velocity at cell edge before and after treatment with either Sema3A_S-P_, Jasplakinolide or mutant Sema3A (n=25-33 cells per group) using same experimental setup as in (B). Data combined from three independent experiments and indicate mean ± SD. **** = P < 0.0001, ns = not significant, by paired t-test. **F.** Intensity plot of LifeAct signal before and after Sema3A_S-P_ treatment of a single OT-I T cell (left) or quantified on multiple cells exposed to Sema3A_S-P_ (right) using same experimental setup as in (B). Arrows indicate measured cell width. ** = P < 0.01, by paired t-test. **G.** Cell width dynamics measured like (F) over time before (white background) or after (grey background) Sema3A_S-P_, Jasplakinolide or mutant Sema3A addition to medium. **H.** Quantification of IRM area of individual OT-I T cells (grey lines) or average for group (red line) over time, with no treatment (leftmost white background), under treatment with Sema3A (grey background) and then Blebbistatin (rightmost white background). Above representative contour plots of single cell under different treatments, with color denoting time (150 sec total). Cells were allowed to settle, and form contact for 3-5 min before data acquisition. Area normalized to cell area at t = 0 sec. Data combined from three independent experiments (n=27 cells). *** = P < 0.001, **** = P < 0.0001, ns = not significant by two-way ANOVA at time-points 90, 270 and 450 sec. **I.** Quantificantion of IRM area of individual OT-I T cells and representative contour plots as in (H), but with treatment with Blebbistatin (grey background) before Sema3A_S-P_ (rightmost white background). Data combined from three independent experiments (n=37 cells). * = P < 0.05, ns = not significant by two-way ANOVA at time-points 90, 270 and 450 sec. Abbreviations: Min, minutes. Sec, seconds. t, time.

Sema3A signaling through Plexin-A1 inactivates the small GTPase Rap1A (*32*), which in turn modulates myosin-IIA activity in diverse cell types (*33, 34*). The effects on the T cell cytoskeleton we observed in the presence of Sema3A_S-P_ appeared consistent with increased myosin-IIA activity. We therefore visualized and quantified the contact area of undulating T cells before and after Sema3A_S-P_ treatment followed by treatment of the myosin-II inhibitor Blebbistatin. As the border of IRM and F-actin signal overlay completely (**Supplementary Figure 3F**), we quantified IRM area to avoid phototoxic effects and inactivation of Blebbistatin, which would be caused by exciting LifeAct (*35*). When Sema3A was added, T cell contact area contracted significantly and cells became immobilized, in line with our analysis of F-actin (**Figure 4F-G**). However, when Blebbistatin was added, T cells started undulating and regained their former size (**Figure 4H, Movie S5**). Conversely, when cells were pre-treated with Blebbistatin followed by Sema3A_S-P_, they retained their shape and activity (**Figure 4I, Movie S6**). We therefore conclude that Sema3A inhibits F-actin dynamics in CD8+ T cells, through hyper-activation of myosin-IIA.

### Nrp1-deficiency enhances anti-tumor activity of CD8+ T cells against Sema3A-rich tumors

To investigate the functional importance of Sema3A in suppressing T cell migration and IS formation *in vivo*, we pursued two complementary lines of enquiry. During development, CD8+ T cells express CD4 molecules during a CD4-CD8 double positive stage (*36*). Thus we crossed LoxP-flanked (Flox) Nrp1 mice with CD4-Cre mice to generate CD4-Cre X Nrp1^+/+^, CD4-Cre X Nrp1^Flox/+^ and CD4-Cre X Nrp1^Flox/Flox^ mice (hereafter referred to as Nrp1^+/+^, Nrp1^Flox/+^ and Nrp1^Flox/Flox^, respectively), to generate Nrp1-deficient T cells. Disruption of Nrp1 expression on stimulated CD8+ T cells was confirmed by flow cytometric analysis (**Supplementary Figure 4A**), thereby generating mice with T cells insensitive to Sema3A ligation. Mice bred normally, had no gross anatomical differences, grew at similar rates and showed no sign of splenomegaly (**Supplementary Figure 4B-C**). Analysis of thymocyte subsets and differentiated T cell memory subsets in the spleen revealed no differences between genotypes (**Supplementary Figure 4D-E**), suggesting that NRP1 is not involved in thymocyte development or T cell homeostasis in non-inflamed conditions. CD8+ T cells from mice of all genotypes expressed similar levels of effector cytokines following CD3/CD28 stimulation (**Supplementary Figure 4F**). We next set out to establish the role of NRP1 on CD8+ T cell priming and activation by infecting mice with the A/PR/8/34-derived pseudotyped influenza virus H7 (Netherlands/2003) N1 (England/2009) (here called H7N1 S-Flu). This virus is capable of triggering strong H-2 D^b^-restricted influenza nucleoprotein (NP)-specific CD8+ T cell responses, but due to suppression of the hemagglutinin (HA) signal sequence cannot replicate or generate anti-HA specific neutralizing antibodies (*37*). This allowed us to specifically consider T cell responses. Mice were infected intranasally with H7N1 S-flu and weighed daily. No differences in weight between genotypes was observed (**Supplementary Figure 4G**). We detected no differences in percentage or absolute number of H-2 D^b^ NP-tetramer positive CD8+ T cells in lungs, draining lymph nodes (dLN) or spleen, ten days post-infection (**Supplementary Figure 4H-I**). Examining the phenotype of CD8+ T cells in the lung, we found that infected mice from all genotypes had an expansion of effector T cells as compared to uninfected mice (**Supplementary Figure 4J**). Consequently, we conclude that NRP1 is dispensable for CD8+ T-cell priming and activation.

We then challenged Nrp1^+/+^, Nrp1^Flox/+^ and Nrp1^Flox/Flox^ mice with B16.F10 and Lewis lung carcinoma (LL/2) cells. The poor immunogenicity of both cell-lines has been overcome using combination therapies that augment immune responses, such as anti-PD1 and anti-4-1BB (*38, 39*), and we therefore considered them good models for examining T cell anti-tumor activity. Notably, Nrp1^Flox/Flox^ mice had significantly delayed tumor growth and increased survival when challenged with either B16.F10 or LL/2 (**Figure 5A-B, Supplementary Figure 4K**). We confirmed that this effect was dependent on CD8+ T cells, as antibody-mediated depletion of CD8+ T cells allowed B16.F10 cells to grow unperturbed in Nrp1^Flox/Flox^ mice (**Figure 5C, Supplementary Figure 4L**). When examining levels of tumor-infiltrating lymphocytes (TILs) in Nrp1^+/+^, Nrp1^Flox/+^ and Nrp1^Flox/Flox^ mice, we noticed a significant increase in the numbers of CD8+ T cells within tumors in Nrp1^Flox/Flox^ mice, but not of CD4+ T cells (**Figure 5D, Supplementary Figure 4M**). Bone-marrow (BM) chimeric mice, containing mixed Nrp1^Flox/+^ and Nrp1^Flox/Flox^ BM, confirmed that the increased levels of infiltration were intrinsic to CD8+ T cells themselves and not dependent on unspecific effects by other cell types (**Figure 5E**).

**Figure 5:**
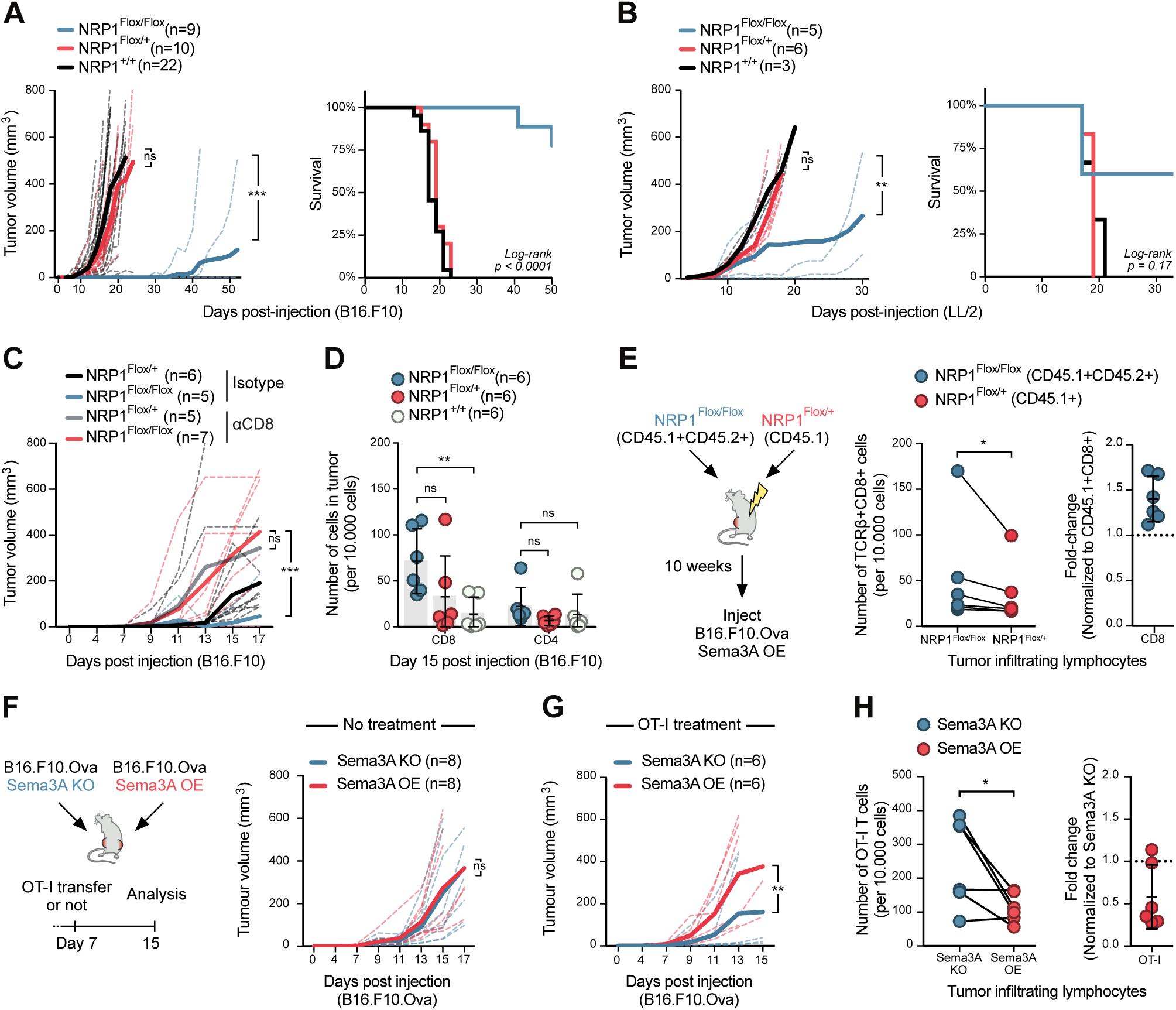
Nrp1-deficiency enhances anti-tumor migration and activity of CD8+ T cells. **A.** Growth curve of B16.F10 cells in NRP1^+/+^, NRP1^Flox/+^ and NRP1^Flox/Flox^ mice (left) and Kaplan-Meier survival curve (right). Dashed lines indicate growth in individual mice, bold line average for group. Combined data from 4 independent experiments with 3-6 mice per group. *** = P < 0.001, ns = not significant by two-way ANOVA. **B.** Growth curve of LL/2 cells in NRP1^+/+^, NRP1^Flox/+^ and NRP1^Flox/Flox^ mice (left) and Kaplan-Meier survival curve (right) (n=3-6 mice per group). Dashed lines indicate growth in individual mice, bold line average for group. Experiment performed once. *** = P < 0.001, ns = not significant by two-way ANOVA. **C.** Growth curve of B16.F10 cells in NRP1^Flox/+^ and NRP1^Flox/Flox^ mice pre-treated with either anti-CD8 antibody or isotype control (n=5-7 mice per group). Dashed lines indicate growth in individual mice, bold line average for group. Data combined from two independent experiments. *** = P < 0.001, ns = not significant by two-way ANOVA. **D.** Enumeration of CD4+ and CD8+ T cells infiltrated into B16.F10 tumors in NRP1^+/+^, NRP1^Flox/+^ and NRP1^Flox/Flox^ mice (n=6 per group). Data indicate mean ± SD. ** = P < 0.001, ns = not significant by two-way ANOVA. **E.** Experimental setup of mixed bone-marrow chimeras in C57BL/6 mice (left) and subsequent enumeration of CD8+ T cells in mice (middle graph). Ratio of CD8+ T cells from NRP1^Flox/Flox^ to NRP1^Flox/+^ bone-marrow derived cells (right graph) (n=6 mice per group). Experiment performed once. Data indicate mean ± SD. * = P < 0.05 by one-way ANOVA. **F.** Experimental setup using B16.F10 Sema3A KO or Sema3A OE cells (left) and growth curve of cells in untreated mice (right) (n=8 mice). Experiment performed once. ns = not significant by two-way ANOVA **G.** Growth curve of B16.F10 Sema3A KO or Sema3A OE cells using similar experimental setup as in (F), but with OT-I treatment at day 7 post-injection (n=6). Experiment performed once. ** = P < 0.001 by two-way ANOVA. **H.** Enumeration of OT-I T cells in tumors (left graph) and their ratio of cells, normalized to the number in the B16.F10 Sema3A KO tumor (right) from same experiment as in (G). * = P < 0.05 by two-way ANOVA.

We hypothesized that the reason T cell immunity was enhanced by NRP1-deficiency in our tumor models, but not against H7N1 S-flu, was an increased availability of Sema3A in the former. Indeed, we did not find high levels of Sema3A on either epithelial cells, leukocytes or endothelial cell-subsets in the lung before, during or after infection with H7N1 S-flu (**Supplementary Figure 5A-B**). Conversely, aggressively growing tumors such as B16.F10, often generate a hypoxic TME (*40*) which itself can induce Sema3A production (*22*). We cultured B16.F10 cells in normoxic or hypoxic conditions and performed RT-qPCR. As expected, hypoxic conditions led to upregulation of known hypoxic response genes, including Pdk1, Bnip3 and Vegfa, in addition to upregulation of Sema3A transcript (**Supplementary Figure 5C**). Flow cytometric analysis of B16.F10 cells grown for 11 days *in vivo* confirmed expression of Sema3A within the TME (**Supplementary Figure 5D-E**). In order to better control the level of Sema3A within the TME, we therefore generated B16.F10.Ova cells that either overexpress or lack Sema3A, upon gene disruption by CRISPR/Cas9 (here referred to as Sema3A OE and Sema3A KO, respectively). Deep-sequencing, RT-qPCR for Sema3a transcript, and analysis by flow cytometry, confirmed that cells lacked or over-expressed Sema3A (**Supplementary Figure 5F-H**). Sema3A OE and Sema3A KO cell-lines grew at similar rates compared to wild-type B16.F10.Ova cells under both normal growth conditions and in the presence of the proinflammatory cytokines IFNγ and TNFα *in vitro* (**Supplementary Figure 5I**). Importantly, when we injected Sema3A OE and KO cell lines into opposite flanks of C57BL/6 mice, tumors grew at similar rates (**Figure 5F**), thus confirming that the cell-lines had a comparable phenotype and growth potential. However, when we adoptively transferred stimulated OT-I T cells into these mice, Sema3A KO tumor growth was significantly delayed compared to Sema3A OE tumors (**Figure 5G**). These results demonstrate that Sema3A overexpression within the TME was sufficient to effectively suppress tumor-specific killing. Furthermore, significantly fewer OT-I T cells had infiltrated tumors that overexpressed Sema3A, compared to Sema3A KO tumors (**Figure 5H**). Taken together, our data underscores the functional significance of Sema3A within the TME as a potent inhibitor of CD8+ T cell migration, and thereby anti-tumor immunity, via interaction with NRP1.

### NRP1 is expressed on tumor-infiltrating CD8+ T cells in clear cell renal cell carcinoma patients

We wished to explore if our findings were relevant to human cancer. Analysis of publicly available TCGA data revealed that high Sema3A expression was associated with poorer survival in clear cell renal cell carcinoma (ccRCC) (**Figure 6A**). We hence turned to a cohort of ccRCC patients that had undergone nephrectomy (**Supplementary Table 1**) to explore the role of the Sema3A/NRP1 pathway in cancer immunity (**Figure 6B**). We first quantified NRP1 expression on CD8+ T cells from peripheral blood (PBMCs), and CD8+ TILs within tumor and tumor-adjacent tissue. Significantly more CD8+ T cells in both tumor and tumor-adjacent tissue expressed NRP1 (**Figure 6C-D, Supplementary Figure 6A**), suggesting that these cells would be sensitive to Sema3A. In our murine model, NRP1 expression correlated with antigen exposure (**Figure 1C**), and we therefore speculated that NRP1-positive CD8+ TILs might be tumor-specific. Indeed, most NRP1+ TILs were also PD1-positive (**Figure 6E-F**), demonstrating that they had either recently been activated or experienced chronic exposure to antigen (*41*). To investigate this, we single-cell sorted NRP1-negative and positive CD8+ T cells from four ccRCC patients and examined their TCR repertoire. TCR diversity, as calculated by either Shannon (SA) and Simpson (SI) diversity indices of CDR3β (**Figure 6G**) and TRBV usage (**Figure 6H**), showed that NRP1+ CD8+ TILs were more clonal than NRP1-T cells, further supporting the hypothesis that such T cells had undergone clonal expansion following recognition of their cognate antigens (*42*) (**Figure 6E-F**). Various cancer-testis (CT) antigens can be expressed by cancerous tissue in ccRCC (*43*). We took advantage of this fact and screened four HLA-A2-positive patients for the presence of HLA-A2-restricted CT-antigen-specific CD8+ T cells using a panel of 21 HLA-A2 tetramers loaded with CT epitopes (*44*). In the three patients, who had CT tetramer positive TILs, we found that a larger proportion of NRP1+ CD8+ TILs were specific for CT-antigens (**Figure 6I-J, Supplementary Figure 6B**). Taken together these data show that NRP1+ CD8+ TILs were found in ccRCC patients, were activated, and were likely specific for tumor-associated antigens.

**Figure 6:**
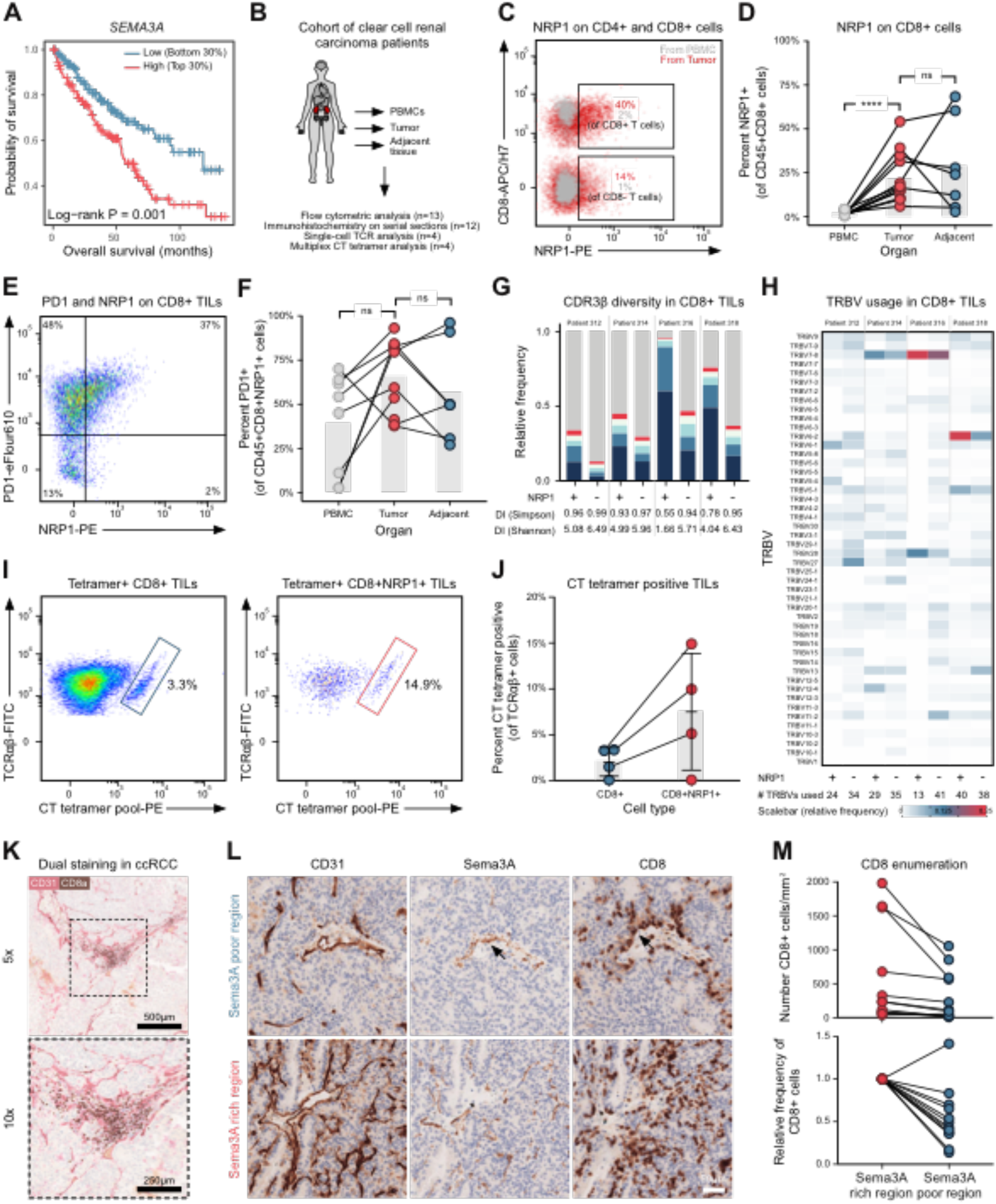
CD8+ TILs express NRP1 and are captured in Sema3A rich areas in ccRCC tumors. **A.** Correlation of SEMA3A mRNA level with survival of ccRCC patients. Data from TCGA, using TIMER (*71*). **B.** Schematic representing ccRCC cohort of patient utilized in (C-M). **C.** Representative flow cytometric analysis of CD8 and NRP1 expression in PBMC and TILs in ccRCC patient. **D.** Analysis of NRP1 expression on CD8+ T cells in PBMC, tumor and tumor-adjacent tissue in ccRCC cohort (n=7-13). Bars indicate mean. **** = P < 0.0001, ns = not significant by two-way ANOVA. **E.** Representative flow cytometric analysis of PD1 and NRP1 on CD8+ TIL in ccRCC patient. **F.** Analysis of PD1 expression on NRP1 positive CD8+ T cells in PBMC, tumor and tumor-adjacent tissue in ccRCC cohort (n=7-13). Bars indicate mean. ns = not significant by two-way ANOVA. **G.** Analysis of CDR3β diversity in NRP1 positive (+) and negative (-) CD8+ TILs (n=4). Colored bars represent the five most abundant clonotypes. Grey bar represents remaining sequences. SI and SA diversity indices (DI) show that in all four patients, NRP1+ TILs are less diverse. **H.** Heatmap of TRBV usage in NRP1 positive (+) and negative (-) CD8+ TILs (n=4). Color indicates relative usage within all of individual patients, as indicated by scalebar. **I.** Representative flow cytometric analysis of TCRαβ and CT tetramer positive CD8+ TILs (left) and NRP1+ CD8+ TILs (right). Error bars indicate mean ± SD. **J.** Graph of percentage CT tetramer positive NRP1+ (red) and NRP1-(blue) CD8+ TILs in four ccRCC patients. **K.** Representative CD8 (brown) and CD31 (red) staining in ccRCC tumor. Dashed area in top image indicates zoom area in bottom image. Scalebar, 500μm and 250μm. **L.** Representative CD31, Sema3A and CD8 staining in Sema3A poor region (top row) and Sema3A rich region (bottom row). Arrows indicate association between Sema3A and CD8 staining. Scalebar, 50 μm. **M.** Enumeration of CD8+ TILs in Sema3A rich (red dots) and poor (blue dots) in patients (n=12). Abbreviations: CT, cancer testis. DI, diversity indices. SA, Shannon index. SI, Simpson index. TIL, tumor-infiltrating lymphocytes. TCGA, The Cancer Genome Atlas. TIMER, Tumor Immune Estimation Resource. TRBV, TCR beta chain variable.

We next wished to explore the spatial distribution of Sema3A and CD8+ T cells within the TME. For this purpose, we stained ccRCC tissue sections from 12 patients from our ccRCC cohort for Sema3A by immunohistochemistry (IHC). We observed widespread expression of Sema3A both within the tumor as well as in adjacent non-neoplastic kidney tissue. In the tumor, Sema3A was predominantly expressed by smooth muscle cells within the tunica media of tumor vasculature but also in areas of fibromuscular stroma. In the adjacent tissue, glomerular mesangial cells and smooth muscle cells within peritubular capillaries stained positive for Sema3A (**Supplementary Figure 6C**). Next, strict serial sections from the same formalin-fixed paraffin embedded tissue blocks were stained for CD31 and CD8 and computationally aligned to the Sema3A sections. Pathological review confirmed that expression of Sema3A co-localized with that of the blood vessel marker CD31. Furthermore, CD8+ cells were often located within regions of high Sema3A expression (**Supplementary Figure 6D**); indeed dual-staining of CD31 and CD8 in ccRCC clearly showed that CD8+ cells were restricted to the immediate area surrounding blood vessels (**Figure 6K**). To further explore the effect of Sema3A on CD8+ cell infiltration and localization, we compared regions within each tumor that were either Sema3A-rich or Sema3A-poor, allowing us to control for variability in CD8+ cell infiltration between patients (**Figure 6L**). This analysis confirmed that our selected Sema3A-rich regions expressed more CD31 than the Sema3A-poor regions, underscoring the close association of Sema3A with the vasculature (**Supplementary Figure 6E-F**). In 11 out of 12 examined patients, there were significantly more CD8+ TILs in the Sema3A-rich areas than in Sema3A-poor areas, corresponding to 46% fewer CD8+ cells in Sema3A-poor regions (**Figure 6M**). Additionally, the CD8+ cells that were present in Sema3A-poor regions were often found clustered near sources of Sema3A (**Figure 6L, arrows**). The data presented here are consistent with a role for Sema3A in modulating T cell infiltration and restricting CD8+ cells to perivascular areas within the tumor.

## DISCUSSION

In this study we showed that the secreted protein Sema3A has a previously underappreciated role in controlling tumor-specific CD8+ T cells and highlight several important conclusions concerning its function.

We established several lines of evidence that reveal a strong inhibitory effect of Sema3A on CD8+ T cell migration in tumors. First, *in vitro* experiments provided functional insights into how Sema3A inhibited key steps in T cell extravasation, including adhesion, transmigration and mobility. Notably, these effects could be reversed using a blocking antibody against NRP1, confirming that NRP1 is an important regulator of Sema3A signaling on CD8+ T cells. Second, conditional knockout of NRP1 on T cells corroborated these findings *in vivo*, resulting in higher CD8+ T cell infiltration into the TME. Conversely, significantly fewer tumor-specific T cells homed to and infiltrated Sema3A-overexpressing tumors. Third, in ccRCC patients CD8+ TILs were preferentially found in Sema3A-rich regions and beside Sema3A-rich blood vessels, reminiscent of how tumor-associated macrophages can be entrapped within Sema3A-rich hypoxic regions (*22*).

We explored the effect of Sema3A on IS formation. Previous studies have characterized Sema3A as an inhibitor of T cell signaling and proliferation using *in vitro* assays (*8, 12*). We extended these results and confirmed that key steps in synapse formation are affected, including cell-cell binding, formation of close contact zones and organization of distinct supramolecular activation clusters. These findings are in line with work by Ueda et al who found that Sema3E inhibited IS formation in thymocytes (*27*). We further show that the F-actin cytoskeleton becomes activated following Sema3A exposure. Although further experiments are warranted to draw firm conclusions, this effect is ostensibly dependent on myosin-IIA activity, since we could rescue T cell undulation using the drug Blebbistatin which specifically prevents intra-cellular force generation by myosin-II (*45*). High resolution 3D imaging has shown that myosin-IIA forms bona fide arcs above the pSMAC (*46, 47*) but then moves inwards and contracts, thereby pinching the T cell away during termination of the IS (*24*). This isotropic contraction of the actomyosin arc appears similar to myosin’s role during cytokinesis (*48*). Our data suggest that Sema3A leads to hyperactivation of myosin-II, thus enforcing IS termination. Data does exist to provide a link between Sema3A binding and myosin-II. Biochemical and crystallographic studies have shown that Sema3A signaling converts the small GTPase Rap1A from its active GTP-bound state, to its inactive GDP-bound state following binding to Plexin-A (*15, 32*). In epithelial and endothelial cells, active Rap1-GTP can act as a negative regulator of myosin-II (*33*). It is therefore tempting to speculate that Sema3A, by inhibiting Rap1-GTP activity, leads to hyperactivation of myosin-II. Indeed, in both neurons (*7, 49*) and DCs (*23*), Sema3A has been shown to increase myosin-II activity in line with this interpretation, however much of this pathway needs to be further elucidated in T cells. We propose that Sema3A induces a cellular “paralysis” based on integrin-actinomyosin contraction leading to motility paralysis and immunological synapse preemption.

There is a growing interest in NRP1 in the context of T cell anti-tumor immunity. Much research has focused on regulatory T cells (Treg), as NRP1 can be used to identify thymus-derived regulatory T cells (*50*) and has been shown to play an important role in controlling Treg function and survival (*51*). It has become clear that NRP1 is expressed on dysfunctional tumor-specific CD8+ T cells (*9–11*), indicating that the protein might play an important role in regulating CD8+ T cells as well. We show that initial NRP1 expression is controlled by the level of TCR-engagement, that the protein remains expressed on tumor-specific CD8+ T cells *in vivo* and that NRP1 is found on a subset of tumor-specific CD8+ T cells in human ccRCC patients. These latter results are in line with reports by Jackson et al. (*52*) and Leclerc et al. (*11*) who found that approximately 10% of CD8+ TILs from melanoma patients and 14% of non-small-cell lung cancer patients, respectively, were NRP1 positive. Unlike us, Jackson et al. did not find any role for NRP1 in regulating CD8+ T cells when mice were challenged with a leukemia cell line. An explanation for this discrepancy could be a lack of Sema3A expression in this model. Indeed, we did not find any functional differences between NRP1 knockout and wild-type T cells when challenging mice with H7N1 S-Flu, a pathogen that did not lead to any meaningful upregulation of Sema3A in the lung. Conversely, Sema3A knockout or overexpression in B16.F10.Ova cells was shown to have significant effects on T cell migration and control of tumor growth when treated with tumor-specific CD8+ T cells. An even stronger effect was seen in the lack of tumor growth in our Nrp1^Flox/Flox^ mice. These results are in line with Delgoffe et al. (*51*) and Leclerc et al. (*11*) who show similar control of tumor growth when treating mice with a blocking anti-NRP1 antibody. Hansen et al. also found strong anti-tumor effects in a comparable conditional NRP1 knockout model but ascribed this to decreased Treg infiltration into the TME (*53*). More likely, the remarkable control of tumor growth seen by us and others is due to synergistic effects, as our data would suggest that ablation of NRP1 enhances CD8+ T cell migration and effector functions as well. Research by Vignali and colleagues has shown that NRP1 plays a key role in Treg survival and suppressive capabilities within the TME through ligation with Sema4A (*51, 54*). Why does NRP1 enhance Treg function, but inhibit CD8+ T cells? While not exploring this question in detail, we did find that Tregs to a larger extent expressed other NRP1 co-receptors, including TGFβRI and II. Indeed, NRP1 has been shown to enhance TGFβ binding in Tregs (*18*). As Tregs are dependent on TGFβ for their function (*55*), one intriguing possibility is that NRP1 preferentially partners with these TGFβ-receptors on Tregs, while the only co-receptors available on CD8+ T cells are the proteins of the Plexin-A family, which could provide a molecular basis for distinct signaling in each cell type.

Our study highlights an underappreciated tumor-escape mechanism, namely inhibition of tumor-specific T cells through cytoskeletal paralysis. We find that the effects of Sema3A on CD8+ T cells are mainly mediated through the co-receptor NRP1, suggesting new therapeutic avenues, for example by using antagonistic NRP1 antibodies. Enhancing migration of tumor-specific T cells into tumors is critical for improving the efficacy of checkpoint blockade (*56*) and adoptive T cell transfer therapies (*57*), making this an exciting prospect. However, since the Sema3A-Plexin-A-NRP1 pathway also regulates the maturation of endothelial cells (*15*) emphasis on timing and drug-target will be critical.

## METHODS

### Cell lines and media

Cell culture was performed using antiseptic techniques in HEPA filtered culture cabinets. Cell lines were grown at 37c in a 5% CO_2_ atmosphere. As indicated in text, for some experiments, cells were cultured for 24 hours in a 1% O_2_ chamber. All cell-lines were screened for Mycoplasma. Adherent cells where split by Trypsin-EDTA detachment and serially passaged and their viability regularly checked.

B16.F10 and B16.F10.Ova cell-lines were provided by Uzi Gileadi. The latter was generated by transducing B16.F10 with a modified Ovalbumin construct, containing a start codon and amino-acid 47 to 386 of the full-length ovalbumin, which ensures that Ovalbumin is not secreted by the cell-line. LL/2 cells were a gift from Christopher W Pugh (Nuffield Department of Medicine, University of Oxford).

B16.F10.Ova Sema3A knockout cells were generated using CRISPR/Cas9 genome-editing (see below). B16.F10.Ova Sema3A overexpressing cells were generated by transducing cells with a lentivirus encoding EFS-Sema3A cDNA (NCBI sequence NM_001243072.1)-mCherry, cloned by VectorBuilder (see below). HEK293T cells were a gift from Tudor A. Fulga (Radcliffe Department of Medicine, University of Oxford).

Adherent cells were kept in DMEM, 10% FCS, 2 mM Glutamine, 1 mM Sodium Pyruvate, 100 U/ml penicillin + 100 μg/ml streptomycin. For some experiments, 10 ng/mL murine IFNγ (cat. no 315-05, PeproTech) or murine TNFα (cat. no. 315-01A, PeproTech) was added to medium. T cells were kept in IMDM, 10% FCS, 2 mM Glutamine, 1 mM Sodium Pyruvate, 1x Non-essential amino acids, 100 U/ml penicillin + 100 μg/ml streptomycin, 10 mM HEPES, 50 μM β-mercaptoethanol. 10 IU IL-2 (cat. no AF-212-12, PeproTech) was added from frozen stock just before use.

### Mouse strains and injection of tumor cells, T cells, antibodies and S-Flu H7N1

All experiments were performed in mice on a C56BL/6 background. Mice were sex-matched and aged between 6 and 12 weeks at the time of the first experimental procedure. All studies were carried out in accordance with Animals (Scientific Procedures) Act 1986, and the University of Oxford Animal Welfare and Ethical review Body (AWERB) under project licence 40/3636. CD4-Cre mice were a gift from Katja Simon (NDM, University of Oxford). LifeAct mice were a gift from Shankar Srinivas (DPAG, University of Oxford). C57BL/6, OT-I and CD45.1 mice were purchased from Biomedical services, University of Oxford. NRP1-floxed mice were purchased from Jackson Laboratories (Stock No: 005247).

Cancer cell lines were split at 1:3 ratio 24 hours before injection into mice in order to keep cells in log-phase. On the day of injection, cells where trypsinized and washed 3 times in PBS to remove residual FBS. Suspensions of 1.5 × 10^5^ cells in 100 uL PBS were prepared and kept on ice until injection. Mice were anesthetized using isoflurane and cells injected intradermally.

For adoptive transfer of T cells into mice, OT-I splenocytes were stimulated for 48 hours using SIINFEKL peptide and sorted as described below, washed 2 times in PBS and injected i.v. via the tail vein.

For infection with S-Flu H7N1, mice were infected intranasally with 10 infectious units S-Flu H7N1 in 50uL viral growth medium (DMEM with 2 mM Glutamine, 10 mM HEPES, 100 U/ml penicillin + 100 μg/ml streptomycin and 0.1% BSA) under anesthesia.

For CD8-depletion experiments, anti-CD8a (cat. no. BE0061, clone 2.43, BioXcell) or IgG2b isotype control (cat. no. BE0090, clone LTF-2, BioXcell) were resuspended in PBS and injected intraperitoneally at day -4, -1, 4 and 7 post injection of cancer cells.

### Mixed bone-marrow chimeras

To generate mixed bone marrow chimeric mice, male C57BL/6 host mice were lethally irradiated at 4.5 Gy for 300 seconds, followed by a 3 hour rest, and a subsequent 4.5 Gy dose for 300 seconds. Mice were injected i.v. with a 1:1 mixture of CD45.1+ NRP1^Flox/+^ and CD45.1+CD45.2+ NRP1^Flox/Flox^ bone marrow cells. Recipient mice received drinking water containing antibiotics (0.16mg/mL Enrofloxacin (Baytril), Bayer Coporation). Mice were rested for 10 weeks before experimental use.

### Analysis of publicly available transcriptional data

For analysis of Sema3A co-receptors, we downloaded raw expression data collected from mice from the “Immunological Genome Project data Phase 1” via the Gene Expression Omnibus (series accession: GSE15907). We specifically focused on naïve CD8+ T cells (accessions: GSM605909, GSM605910, GSM605911), CD8+ effector T cells (accessions: GSM538386, GSM538387, GSM538388, GSM538392, GSM538393, GSM538394), and CD8+ memory T cells (accessions: GSM538403, GSM538404, GSM538405). The raw expression array files were processed using the affy package (*58*) and differential expression of selected genes (CD72, NRP1, NRP2, PLXNA1, PLXNA2, PLXNA3, PLXNA4, PLXNB1, PLXNB2, PLXNB3, PLXNC1, PLXND1, SEMA3A, SEMA3B, SEMA3C, SEMA3D, SEMA3E, SEMA3F, SEMA3G, SEMA4A, SEMA4B, SEMA3C, SEMA4D, SEMA4F, SEMA4G, SEMA5A, SEMA5B, SEMA6A, SEMA6B, SEMA6C, SEMA6D, SEMA7A, TIMD2, HPRT, OAZ1, RPS18, NFATC2, TBX21, EOMES, CD28, PDCD1, CTLA4, LAG3, BTLA, TIM3, ICOS, TNFRSF14, TNFSF14, CD160, CD80, LAIR1, CD244, CXCR1, CXCR2, CXCR3, CXCR4, CXCR5, CCR1, CCR2, CCR3, CCR4, CCR5, CCR5, CCR6, CCR7, CCR8, CCR9, CCR10) between naïve and effector and memory cells was examined using the limma package (*59*) in R (*60*).

Analysis of TCGA data was conducted using TIMER (*61*).

### Harvesting and activating splenocytes

Mice were euthanized using CO_2_, and spleens were harvested and stored in T cell media on ice. The spleen was strained through a 70 μm nylon mesh using the blunt end of a syringe to make a single cell suspension. Cells were washed off the mesh by applying 5 mL of T cell medium, followed by mixing of the solution by aspiration. Cells were then washed and resuspended in 3 mL red blood-cell (RBC) lysis buffer for 5 minutes on ice. Cells were washed again in T cell medium, counted and resuspended at 2×10^6^ cells per mL in T cell medium.

10 IU/ml IL-2 (Cat. 212-12, PeproTech) and 25nM SIINFEKL (N4) peptide (Cambridge Peptides) were added to the single cell solution. Approximately 200 μL cells were then plated onto a 96-well U-bottom plate and allowed to expand for 48 hours. For TCR affinity assays, SIINFEKL (N4), SIITFEKL (T4) or SIIQFEKL (Q4) peptide (Cambridge Peptides) was used at indicated concentrations.

### Sorting CD8+ T cells using magnetic beads

CD8a+ Negative T Cell Isolation Kit (Order no. 130-104-075, Miltenyi Biotec) was used to sort T cells and was performed according to the manufacturer’s protocol. Briefly, cells were washed in MACS buffer (0.5% bovine serum albumin and 2 mM EDTA in PBS), incubated with antibody cocktail, followed by magnetic beads for 10 minutes each on ice. Cells were then loaded into a prewetted LS column (Order no. 130-042-401, Miltenyi Biotec) inserted into a magnet in approximately 3-5 mL MACS buffer.

### Preparation of tissue from mice for flow cytometry

When staining cells in B16.F10 and LL/2 tumors, or from lymph nodes, frontal cortex, lungs or thymus, mice were euthanized using CO2, and tumors were harvested and stored in T cell media on ice. Organs were cut into smaller pieces with a scalpel and incubated for 30 minutes with reagents from a tumour dissociation kit (Order no. 130-096-730, Miltenyi Biotec). Cells were strained through a 70 μm nylon mesh using the blunt end of a syringe to make a single cell suspension. Cells were washed off the mesh by applying 5 mL T cell media, followed by mixing of the solution by aspiration. After a wash, the cells were resuspended in approximately 2 mL of 100% Percoll solution (Cat. no. 17-0891-01, GE Healthcare), and layered carefully on top of 3 mL of 80% and 40% Percoll solution, and spun for 30 minutes at 2000g. Cells at the 80-40% interphase were washed, and stained using protocols outlined below.

### Flow cytometry

For washing and staining cells for flow cytometry PBS with 2% BSA, 0.1% NaN3 sodium azide was used. Single colour controls were either cells or OneComp Compensation Beads (Cat. No 01-1111-41, Thermo Fisher).

For surface staining, cells were washed with 200 μl FACS Buffer and blocked in 100 μl Fc block (cat. no. 101319, TruStain FcX, clone 93, BioLegend, diluted 1:100) in FACS Buffer for 10 min on ice and washed. Antibody cocktail was added and cells were stained on ice for at least 20 min, in the dark and washed twice. When applicable, cells were fixed in 2% PFA for at least 10 min at RT before acquisition. For quantification of number of cells in tumors, lymph nodes and lungs in certain experiments, quantification beads (CountBright, cat. no. C36950, Thermo Fisher) were used.

For intracellular staining, cells were fixed in 100 μl/well of FoxP3 IC Perm/fix Buffer (Cat. no 00-8222-49, Thermo Fisher) for 20 min at RT. Cells were pelleted, the fixative removed, and 200 μl/well of 1x Perm Buffer added for 20 min at RT and washed. Antibody mix was added and cells stained for at least 20 min on ice in Perm buffer and washed twice.

Cells were analyzed on either Attune NxT (Life Technologies) or an LSR Fortessa X20 or X50 (BD Biosciences) flow cytometers in the WIMM Flow Cytometry Facility and data was analysed using FlowJo v10 (FlowJo) and R (*60*).

The following antibodies and tetramers were used for flow cytometry:

| Specific for | Fluorophore(s) | Company | Clone |
| --- | --- | --- | --- |
| CD3e | BV650 | BioLegend | 17A2 |
| CD4 | APC710 | Tonbo<br>Bioscience | GK1.5 |
| CD4 | BUV810 | BD | GK1.5 |
| CD8a | BV711 | BioLegend | 53-6.7 |
| CD11a | PE | BioLegend | M17/4 |
| CD11b | APC | BioLegend | M1/70 |
| CD19 | BV425 | BioLegend | 6D5 |
| CD24 | APC710 | BD | M1/69 |
| CD25 | PerCP/Cy5.5 | BioLegend | 3C7 |
| CD31 | BV510 | BD | MEC13.3 |
| CD44 | APC/Cy7,<br>PE/Cy7 | BioLegend | IM7 |
| CD45.1 | FITC | eBioscience | A20 |
| CD45.2 | PerCP/Cy5.5 | BioLegend | 104 |
| CD49d | BUV395 | BD | 9C10 |
| CD62L | BV610,FITC | BioLegend | Mel-14 |
| CD69 | BUV737 | BD | H1.2F3 |
| CD103 | PE/Cy7 | BioLegend | 2E7 |
| CD105 | PE-CF594 | BD | MJ7/18 |
| CD162 | BV421 | BD | 2PH1 |
| EpCAM | APC/Cy7 | BioLegend | G8.8 |
| F4/80 | BV610 | BioLegend | BM8 |
| FoxP3 | BV421 | BioLegend | MF-14 |
| IFNγ | PE | BioLegend | XMG1.2 |
| Granzyme B | FITC | BioLegend | GB11 |
| Ly6C | BV780 | BioLegend | HK1.4 |
| MHC-II | PerCP/Cy5.5 | BioLegend | AF6-120.1 |
| NRP1 | BV421 | BioLegend | 3E12 |
| NRP1 | PE | BioLegend | 3E12 |
| H-2D<sup>B</sup>-NP<br>tetramer | PE-Cy7 | Made in-house |  |
| PlexinA1 | PE | R&D Systems | FAB4309P |
| PlexinA2 | APC | R&D Systems | FAB5486A |
| PlexinA4 | Conjugated to<br>PE | Abcam | ab39350 |
| Podoplanin | APC | BioLegend | 8.1.1 |
| TCRβ | APC/Cy7 | BioLegend | H57-597 |
| TCRβ | PE-CF594 | BD | H57-597 |
| TGFβRI | PE | R&D Systems | FAB5871P |
| TGFβRII | PE | R&D Systems | FAB532P |
| TNFα | PerCP/Cy5.5 | BioLegend | MP6-XT22 |
| Sema3A | FITC, PE | R&D Systems | IC1250P |
| VEGFR2 | PE | BioLegend | Avas12 |
| Zombie | Violet Dye | BV421 | BioLegend |
| Zombie | Aqua Dye | BV525 | BioLegend |
| Zombie | Near-infrared | APC-780 | BioLegend |

### RT-qPCR

RNA was extracted from cells using RNeasy kit (QIAGEN), followed by quantification on Nanodrop (Thermo Scientific). RNA was reverse transcribed using QuantiTect Reverse Transcription Kit (QIAGEN). Both controls without RNA or reverse transcription were included, and all experiments were performed in minimum technical triplicates and biological duplicates. cDNA was diluted to 10-20 ng in 5 ul/well and added to qRT-PCR plates. Taqman probes were combined with 2x Fast Taqman Master Mix and 5 ul/well added to the cDNA. qRT-PCRs were run on a QuantStudio7 qRT-PCR machine (Life Technologies). Expression was normalized to the house keeping gene HPRT.

The following TagMan probes were used: BNIP3 (Mm01275600-g1), HPRT (Mm03024075-m1), PDK1 (Mm00554300-m1), PDL1 (Mm00452054-m1), SEMA3A (Mm00436469-m1) and VEGFA (Mm00437306-m1).

### Western blot

Cells were washed in PBS and pelleted, before being resuspended in lysis buffer with a EDTA protease inhibitor for at least 30 minutes on ice to extract protein. Cell-debris was removed by centrifugation at 4°C. Supernatant containing protein was collected and quantified using Pierce BCA protein assay using diluted albumin as a standard.

Samples were mixed with Loading Buffer (Life Technologies) and Reducing Agent (Life Technologies) and heated to 95°C for at least 5 minutes. 4-12% Bis-Tris gels and MES SDS Buffer (Life Technologies) were used for proteins with a molecular weight below 200 kDa, while proteins above 200 kDa were blotted on a 3-8% Tris-Acetate gels in MOPS Buffer (Life Technologies). Proteins were separated at 200V for approximately one hour and transferred onto either PVDF or nitrocellulose membrane using the TransBlot Turbo Transfer (BioRad) system. Gels were blocked in 5% BSA/PBS solution (blocking buffer) for at least 30 minutes at RT. Membranes were stained with primary antibody in fresh blocking buffer and incubated at 4C overnight on a shaker, washed five times in PBS with Triton-X (0.1% Tween-20 in PBS) followed by incubation with fluorescent secondary antibodies (LICOR) diluted 1:20000 in blocking buffer for 1 hour on a shaker. Membranes were dried and imaged using the Odyssey Near-Infrared imaging system (LI-COR).

The following antibodies were used: Anti-Neuropilin 1 antibody (Abcam, ab184783), anti-PlexinA1 (R&D Systems, AF4309), anti-PlexinA2 (R&D Systems, AF5486), anti-GAPDH (Santa Cruz, sc-32233), anti-β-Actin (Cell Signaling Technology, 13E5).

### CRISPR/Cas9 editing and verification of B16.F10.Ova cells

A sgRNA targeting Sema3a was cloned into Cas9-2A-EGFP expression vector pX458. This vector was electroporated into 1×10 B16.F10.Ova cells suspended in Solution V (Lonza) using the Amaxa 2B nucleofector (Lonza) with settings P-020. After 48 hours, single cells were sorted using the SH800 cell sorter (SONY) and expanded. Clones were genotyped by high-throughout sequencing. Briefly, the targeted locus was PCR amplified from each clone and subsequently indexed with a unique combination of i5 and i7 adaptor sequences. Indexed amplicons were sequenced on the MiSeqV2 (Illumina) and demultiplexed reads from each clone were compared to the wildtype Sema3a reference sequence using the CRISPResso webtool (*62*).

### Lentiviral transduction of cells

A lentiviral Sema3A overexpression vector was purchased from VectorBuilder. To generate viral particles, this transfer vector was co-tranfected into HEK-293T cells along with the packaging and envelope plasmids pCMV-dR8.91 and pMD2.G using polyethylenimine. Crude viral supernatant was filtered and used to transduced B16.F10.Ova cells. mCherry positive transduced cells were selected by FACS using the SH800 cell sorter (SONY) to create a Sema3A overexpression cell line.

### Protein production of Sema3A and mutant Sema3A

Recombinant mouse Sema3A_S_−P (residues 21–568), Sema3A_S-P-I_ (residues 21–675, without a HIS tag) along with the Nrp1-binding deficient mutant Sema3A (residues 21–568, L353N-P355S), here called mutant Sema3A, were cloned into a pHLsec vector optimized for large scale protein production as described before (*63*). The L353N P355S mutation in mutant Sema3A introduces an N-linked glycan to the Sema3A-Nrp1 interaction site, which is sufficient to block the formation of Sema3A-Nrp1-PlxnA2 signaling complex as described previously (*5*). Additionally, in all proteins, furin sites (R551A and R555A) were removed to prevent Sema3A proteolytic processing and enable more controlled purification and sample homogeneity. Proteins were expressed in HEK293T cells. Proteins were purified from buffer-exchanged medium by immobilized metal-affinity followed by size-exclusion chromatography using a Superdex 200 column (GE).

For some experiments, purified Sema3A protein was labelled with AF647 using Alexa Fluor 647 Antibody Labeling Kit according to protocol (Invitrogen, cat. no A20186) at a F/P rate at 1-2.

### Live-cell imaging of cells for migration studies, LifeAct and IRM quantification

u-Slide I 0.4 Luer (Cat. no. 80172, Ibidi), u-Slide 8 well (Cat. no. 80826, Ibidi) and u-Slide Angiogenesis (Cat. no. 81506, Ibidi) were used for live-cell imaging of T cell motility and adhesion.

Proteins were immobilized on the plates by resuspension in PBS and allowing them to adhere for 2 hours as room-temperature, followed by three washes in PBS with 1-2% BSA. The following proteins and concentrations were used: ICAM-1-Fc (Cat. 553006, BioLegend, at 10 μg/mL), CD3 (clone 145-2C11, BioLegend, at 10ug/mL), CXCL12 (Cat. 578702, BioLegend, at 0.4 μg/mL, BioLegend). Sema3A_S-P_ and mutant Sema3A (described above) were either ligated to surfaces in a similar manner or added to the medium while imaging at 5 μM. Plates were used immediately after preparation.

For experiments, T cells were activated for 48 hours using peptide stimulation, sorted and washed. In cases were αNRP1 or isotype control treatment was applied, T cells were incubated with these antibodies for 15 minutes at 37 °C, before an additional wash. T cell medium without phenol red and IL-2 was prewarmed and used to resuspend cells. Cells were added to plates placed on a stage in environmental chamber set at 37C directly at the microscope. DeltaVision Elite Live cell imaging microscope, Zeiss LSM 780 or 880 confocal microscopes were used with a Zeiss Plan-Neofluar 10x (0.3 NA), 40x (0.6 NA)or 63x (1.3 NA) lens.

In cases where T cells were tested under shear stress, cells were loaded into Hamilton syringes, and installed on a Harvard Apparatus PHD 2000 pump, and connected to chambered slides through 0.4 luer tubes. Flow rates and control of directionality of flow (ie, that it was laminar) were measured using fluorescent beads over set distances to calibrate instrument.

In cases were T cells were treated with drugs while imaging, drugs were first resuspended in T cell medium and carefully added on top of solution while images were being acquired. Care was taken always to take into consideration the correct concentration under growing levels of media. The following drugs were used: Jasplakinolide (Cat. J4580, Sigma, used at 5uM) and Blebbistatin (Cat. 203390, EMD Millipore, used at 100 uM), as well as Sema3A_S-P_ (used at 1-5 μM).

Data was acquired at 1 sec to 1 minute per frame as indicated, and analyzed in Fiji/ImageJ (*64*). For cell tracking and visualization of spiderplots the Trackmate package was used (*65*). Subsequent tracks were analyzed in R (*60*) and visualized using the ggplot2 package (*66*). For cell contours, IRM and LifeAct, data was thresholded and collected in Fiji/ImageJ, then exported to R for analysis and visualization. For quantification of IRM and LifeAct area while adding drugs, movies were edited such that the analyzed frames were equal to the timing indicated in figures (ie to start 3-5 minutes before adding the first drug). In videos with a frame-rate of 1 frame per second, 3 frames around the frame in which drugs was added was removed to avoid blurry or distorted images.

### Transwell chemotaxis assay

Trans-migration was assessed in 24-well transwell plates with 3-μm pore size (Corning Life Sciences). The lower chamber was loaded with 500 μl T cell medium with 10 IU IL-2 and with or without 50ng/mL CXCL10. 10^5^ CD8+ OT-I CD8+ cells were stimulated ans sorted as described above and added in a volume of 100 μl of T cell medium to the upper chamber, in either the presence of a blocking αNRP1 antibody or an isotype control, and 5μM Sema3A_S-P_. As a positive control, effector cells were placed directly into the lower chamber. As a negative control, migration medium alone was placed in the upper chamber. Plates were incubated for 3 hours at 37C in a 5% CO_2_ atmosphere. Thereafter, the Transwell inserts were removed and the contents of the lower compartment were recovered. Cells from the lower chamber were stained and the cells were quantified by flow cytometry.

### Immunological synapses analysis on supported lipid bilayers (SLB)

The concentrations of the ligands used were as follows: 5 μg/ml to achieve 10 molecules/μm2 of biotinylated H2Kb-SIINFEKL, 68 ng/ml to achieve 100 molecules/μm2 of 12x-HIS-tagged CD80 (AF488-labelled), and 122 ng/ml to achieve 200 molecules/μm2 of 12x-His tagged ICAM-1 (AF405-labelled). These concentration were determined based on titrations on bead supported bilayers analyzed by flow cytometry. Sema3A_S-P-I_ (described above) was used, as this protein had no HIS-tag and so could not interfere with binding of other tagged proteins in bilayer.

For live cell imaging, supported lipid bilayer presenting H2Kb-SIINFEKL, CD80, and ICAM-1 were assembled in sticky-Slide VI 0.4 luer (Ibidi) channels. The entire channel was filled with a liposome suspension to form bilayer all along the channel. Live cells on SLBs were imaged using the Olympus FluoView FV1200 confocal microscope that was enclosed in an environment chamber at 37 °C and operated under standard settings. 60x oil immersion objective (1.4 NA) was used with 2x digital zoom for time-lapse imaging at 20 second intervals.

For fixed cell imaging, SLBs presenting H2Kb-SIINFEKL, CD80, and ICAM-1 were assembled in 96-well glass-bottom plates (MGB096-1-2-LG-L, Brooks). 50,000 cells were introduced into the wells at 37C and fixed 10 minutes later by adding 8% PFA in 2x PHEM buffer. After 3 washes with 0.1% BSA in HBS the fixed cells were imaged on the InCell 6000 wide-field fluorescence high-throughput imaging station using a 40x air objective (0.75 NA). The imaging station was programmed to visit specific equivalent locations in each of the desired wells in the 96-well plate.

Analysis of fixed cell images was carried out by the MATALB based TIAM HT package (*26*). The source-code is available on the github repository: https://github.com/uvmayya/TIAM HT.

### Acquisition of tissue from ccRCC patients

Acquisition and analysis of ccRCC samples were approved by Oxfordshire Research Ethics Committee C. After informed written consent was obtained, samples were collected, and store until use by Oxford Radcliffe Biobank (project reference 17/A100 and 16/A075).

### Analysis of CDR3 and TRBV usage in CD8+ ccRCC TILs

cDNA from single cells was obtained using a modified version of the SmartSeq2 protocol (*67*). Briefly, single cells were sorted into plates containing lysis buffer. cDNA was generated by template switch reverse transcription using SMARTScribe Reverve Transcriptase (Clontech), a template-switch oligo and primers designed for the constant regions of Trac and Trbc genes. TCR amplification was achieved by performing two rounds of nested PCR using Phusion High-Fidelity PCR Master Mix (New England Biolabs). During the first PCR priming, indexes were included, to identify each cell. A final PCR was performed to add the Illumina adaptors. TCR libraries were sequenced on Illumina MiSeq using MiSeq Reagent Kit V2 300-cycle (Illumina). FASTQ files were demultiplexed for each cell. Sequences from clones were analysed using MiXCR (*68*). Post analysis was performed using VDJtools (*69*).

### CT-tetramer staining

HLA-A2 monomers were made in-house, and loaded with CT-antigens through UV-directed ligand exchange using published protocols (*70*). Following ligand exchange, all monomers were tetramerized through binding to PE-Streptavidin, washed, and combined to allow for “cocktail” staining of cells. Frozen vials of tumor tissue were thawed, dissociated, and CD45-positive sorted, followed by staining with 0.5 μg of each tetramer in 50 ul for one hour at RT. Otherwise staining proceeded like described previously.

The following CT-antigens were loaded into HLA-A2 monomers:

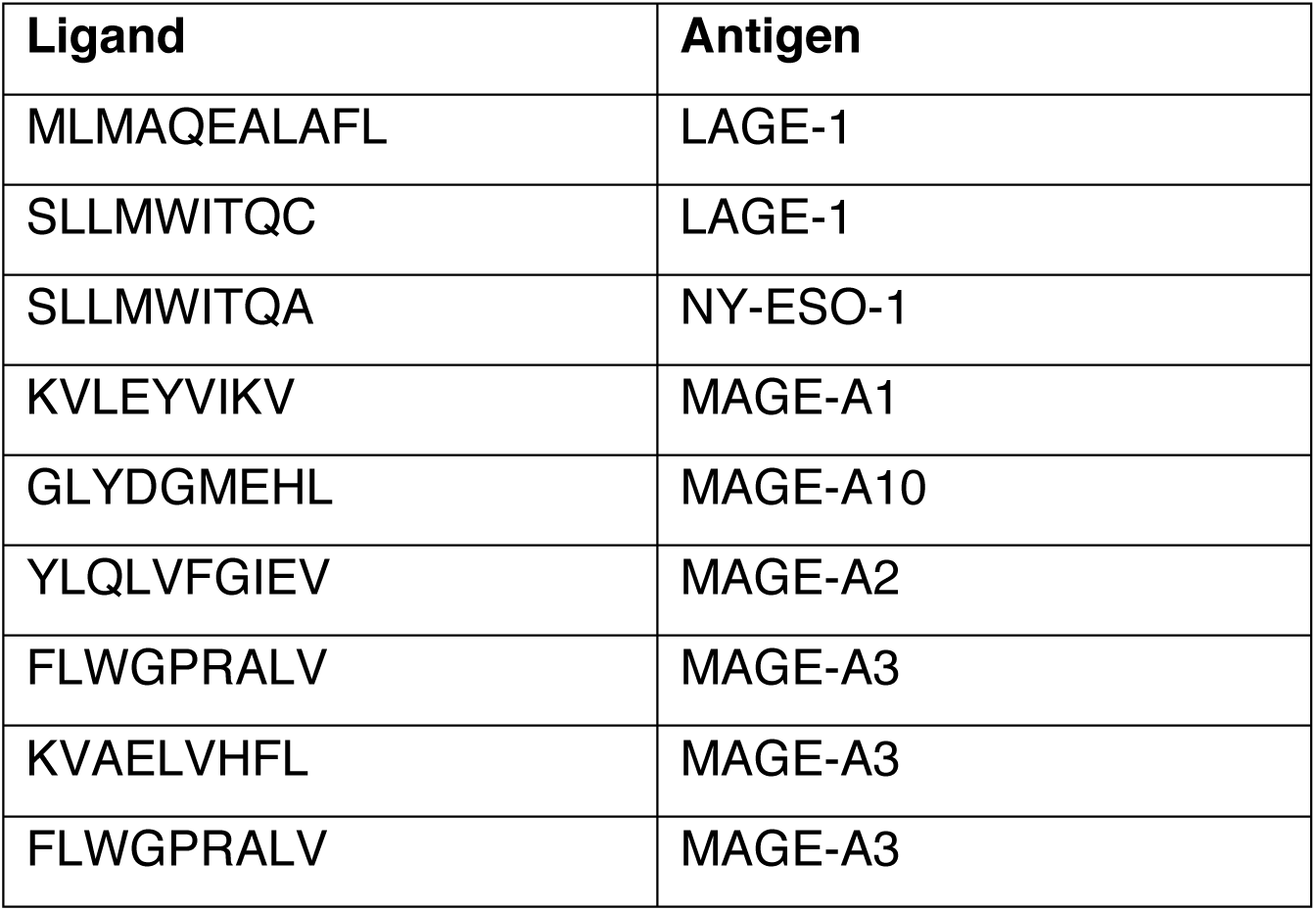
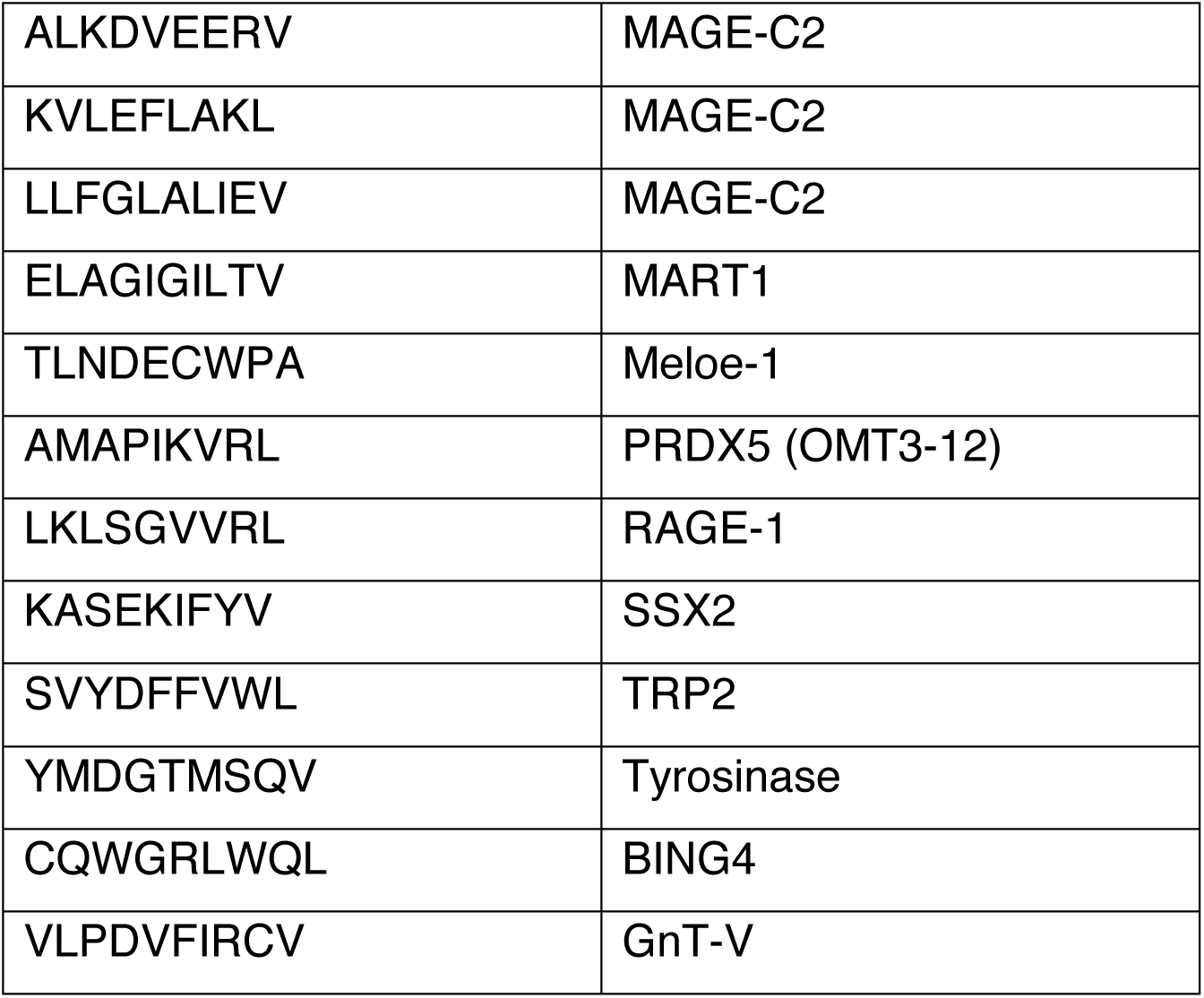

### Immunohistochemistry and image acquisition and analysis

The diagnostic hematoxylin and eosin (H&E) stained slides for 12 cases of clear cell renal cell carcinoma were reviewed to identify corresponding formalin-fixed paraffin embedded tissue blocks that contained both tumor and adjacent non-tumor tissue. Strictly serial 4 μm sections were then cut from the most appropriate block from each case. These sections underwent immunohistochemistry staining on a Leica BOND-MAX automated staining machine (Leica Biosystems). Briefly, sections were deparaffinized, underwent epitope retrieval and endogenous peroxidase activity was blocked with 3% hydrogen peroxide (5 minutes). Subsequently, sections were incubated with the primary antibody (30 minutes) followed by post-primary and polymer reagents (8 minutes each). Next, 3,3’-Diaminobenzidine (DAB) chromogen was applied (10 minutes) (all reagents contained within the BOND Polymer Refine Detection kit, Leica Biosystems, catalogue no. DS9800). For double immunohistochemistry staining, the above cycle was repeated twice with the first cycle using Fast red chromogen labelling (all reagents contained within the BOND Polymer Refine Red Detection kit, Leica Biosystems, catalogue no. DS9390) and the second cycle DAB chromogen labelling. At the end of both the single and double immunohistochemistry protocols, the sections were counterstained with hematoxylin (5 minutes), mounted with a glass coverslip and left to dry overnight. The following primary antibodies were used during staining: CD31 (Agilent Technologies, JC70A, 1:800), CD8 (Agilent Technologies, C8/144B, 1:100) and Sema3A (Abcam, EPR19367, cat. no. ab199475, 1:4000).

Stained slides were scanned at x400 magnification using the NanoZoomer S210 digital slide scanner (Hamamatsu). Sema3A-stained digital images were reviewed by a trained pathologist (PSM) and the extent of staining was quantified in the regions where expression was deemed to be highest (‘Sema3A-rich’) and lowest (Sema3A-poor) within the same tumor, using custom-made Matlab (MathWorks) scripts (% staining = DAB+ pixels/total pixels x 100; raw data, image analysis, and data processing scripts are available upon request). This analysis was repeated in the same regions on adjacent serial sections for CD31 (as for Sema3A) and CD8 (for which discrete cell counts were calculated from stained regions using a water shedding process).

### Statistical analysis

Statistical analysis was performed in Prism software (GraphPad) or R (*60*). Data was tested for Gaussian distribution. For multiple comparisons, either one-way or two-way analysis of variance (ANOVA) was used with Tukey’s test to correct for multiple comparisons. For comparison between two groups, Students t test, Student’s paired t test, or one-tailed or two-tailed Mann–Whitney test were used.

## AUTHORS CONTRIBUTIONS

*Conceptualization*: M.B.B, V.C., E.Y.J., M.L.D., T.A.F., M.F.; *Formal analysis*: M.B.B., Y.S.M., V.A., M.R., P.S.M., V.M., M.F.; *Funding acquisition*: V.C., E.Y.J., M.L.D., T.A.F., M.F., C.W.P.; *Investigation*: M.B.B., Y.S.M., V.A., P.S.M., U.G., S.V., M.R., C.K., J.C., A.V.H., V.M., P.R., L.R.O., M.F., H.C.Y.; *Methodology*: V.C., U.G., M.B.B., M.L.D., M.F., H.C.Y., Y.S.M., A.R.T.; *Project administration*: V.C.; *Resources*: U.G., S.V., V.J., M.R., V.W., Y.K., V.M., A.R.T.; *Software*: M.R., J.A.B., M.B.B.; *Supervision*: V.C.; *Visualization*: M.B.B., P.S.M., V.M., M.F.; *Writing – original draft*: M.B.B., V.C., E.Y.J., M.L.D., M.F.; *Writing – review & editing*: M.B.B., V.C., E.Y.J., M.L.D., M.F., Y.S.M., C.K., P.S.M., V.J.

## ACKNOWLEDGEMENTS

The authors wish to thank members of the Vincenzo Cerundolo and Tudor A. Fulga laboratories, University of Oxford, for helpful discussions and suggestions. Simon Davis. Oliver Bannard and Audrey Gérard, University of Oxford, for helpful advice and guidance. The staff of the University of Oxford Department of Biomedical Services for animal husbandry. Christoffer Lagerholm and Dominic Waithe of the Wolfson Imaging Centre at University of Oxford for microscopy training and support. The Medical Research Council (MRC) WIMM Flow Cytometry Facility for training and support. The Oxford Centre for Histopathology Research and the Oxford Radcliffe Biobank, which are supported by the NIHR Oxford Biomedical Research Centre and the Kennedy Institute of Rheumatology Microscopy Facility. Lisa Browning, Oxford University Hospital, for examination of histology samples. David Pinto, University of Oxford, for code for analysis of CDR3 sequences.

This work was supported by the U.K. MRC (MRC Human Immunology Unit), the Oxford Biomedical Research Centre, and Cancer Research UK (Programme Grant C399/A2291 to V.C.; C375/A17721 to E.Y.J.); the Wellcome Trust (212343/Z/18/Z for M.F.; 100262Z/12/Z for M.L.D. and S.V.) and Kennedy Trust for Rheumatology Research (to M.L.D. and S.V.), European Commission (ERC-2014-AdG_670930 for M.L.D. and V.M.), Cancer Research Institute (to V.M.), and EPSRC (EP/S004459/1 for M.F. and H.C.Y). The Wellcome Centre for Human Genetics is supported by Wellcome Trust Centre grant 203141/Z/16/Z. P.S.M. is supported by a Jean Shanks Foundation/Pathological Society of Great Britain & Ireland Clinical Research Training Fellowship. C.K. is supported by a Wellcome Studentship (105401/Z/14/Z). V.J. is supported by an EMBO Long-Term Fellowship (ALTF 1061–2017). J.A.B. is supported by EPSRC/MRC Centre for Doctoral Training in Systems Approaches to Biomedical Science (EP/G037280/1) and the EPSRC Impact Acceleration Account (EP/R511742/1). L.R.O. is supported by the Independent Research Fund Denmark (8048-00078B).

## SUPPLEMENTARY DATA

**Supplementary Table 1.**

| **Baseline characteristics of patient cohort.** |  |  |
| --- | --- | --- |
| Characteristics | Number<br>(range) | Percent |
| **Age (years)** |  |  |
| Mean | 64.4 (42-86) |  |
| **Gender** |  |  |
| Male | 13 | 56.5% |
| Female | 10 | 43.5% |
| **Tumour grade (ISUP)** |  |  |
| 1 | 0 | 0 |
| 2 | 2 | 8,7% |
| 3 | 12 | 52.2% |
| 4 | 6 | 26.1% |
| N/A | 3 | 13% |
| **Tumour location** |  |  |
| Right | 14 | 60.9% |
| Left | 9 | 39.1% |
| **Type of surgery** |  |  |
| Radical nephrectomy | 17 | 73.9% |
| Partial nephrectomy | 6 | 26.1% |
| **Tumour stage** |  |  |
| pT1a | 1 | 4,5% |
| pT1b | 5 | 22.7% |
| pT2a | 1 | 4.5% |
| pT2b | 0 | 0 |
| pT3a | 14 | 63.6% |
| pT3b | 0 | 0 |
| pT3c | 1 | 4.5% |
| pT4 | 0 | 0 |
| N/A | 1 | 4.5% |

**Supplementary Figure 1 (relates to Figure 1).**
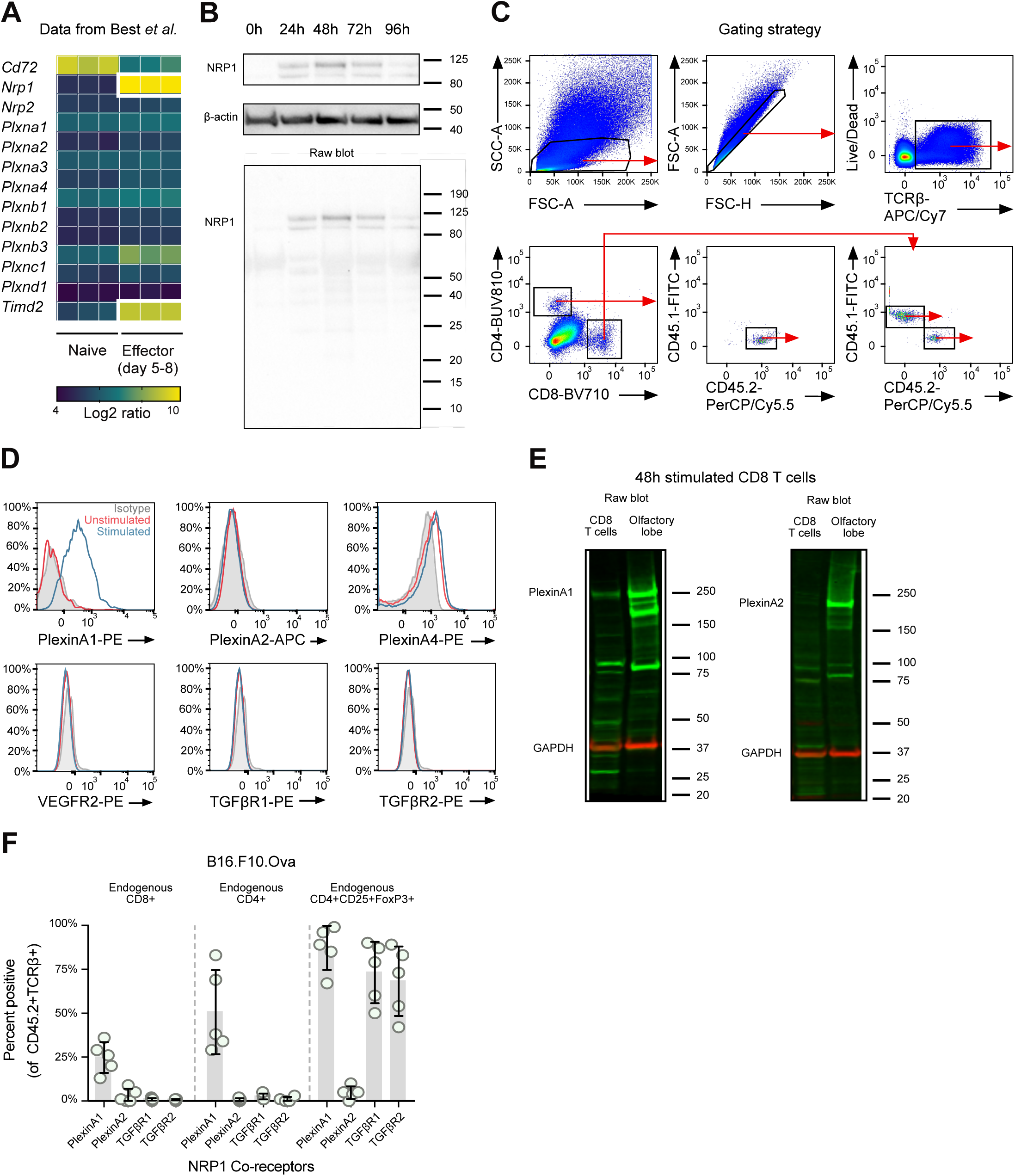
**A.** Heatmap of transcript levels of known semaphorin receptors on naïve and effector OT-I T cells following infection with vaccinia-OVA. Data from Best et al. 2013 (*16*). **B.** Western blot showing NRP1 up-regulation in OT-I T cells following stimulation with SIINFEKL. Experiment was performed once. **C.** Gating strategy for Figure 1F-G. **D.** Flow cytometric analysis of Plexin-A1, Plexin-A2, Plexin-A4, VEGFR2, TGFβR1 and TGFβR2 on unstimulated and 48 hour SIINFEKL stimulated OT-I T cells. Experiment representative of three. **E.** Western blots showing expression of Plexin-A1 (left, green) and Plexin-A2 (right, green) in 48 hour stimulated OT-I T cells and olfactory lobe (positive control). Experiment performed once. Loading control GAPDH shown in red. **F.** Flow cytometric analysis of Plexin-A1, Plexin-A2, TGFβR1 and TGFβR2 expression on OT-I T cells and endogenous CD8+ TILs, 11 days after adoptive transfer og OT-I T cells in antigen-expressing tumor (B16.F10.Ova) (n=5). Error bars indicate SD.

**Supplementary Figure 2 (relates to Figure 2).**
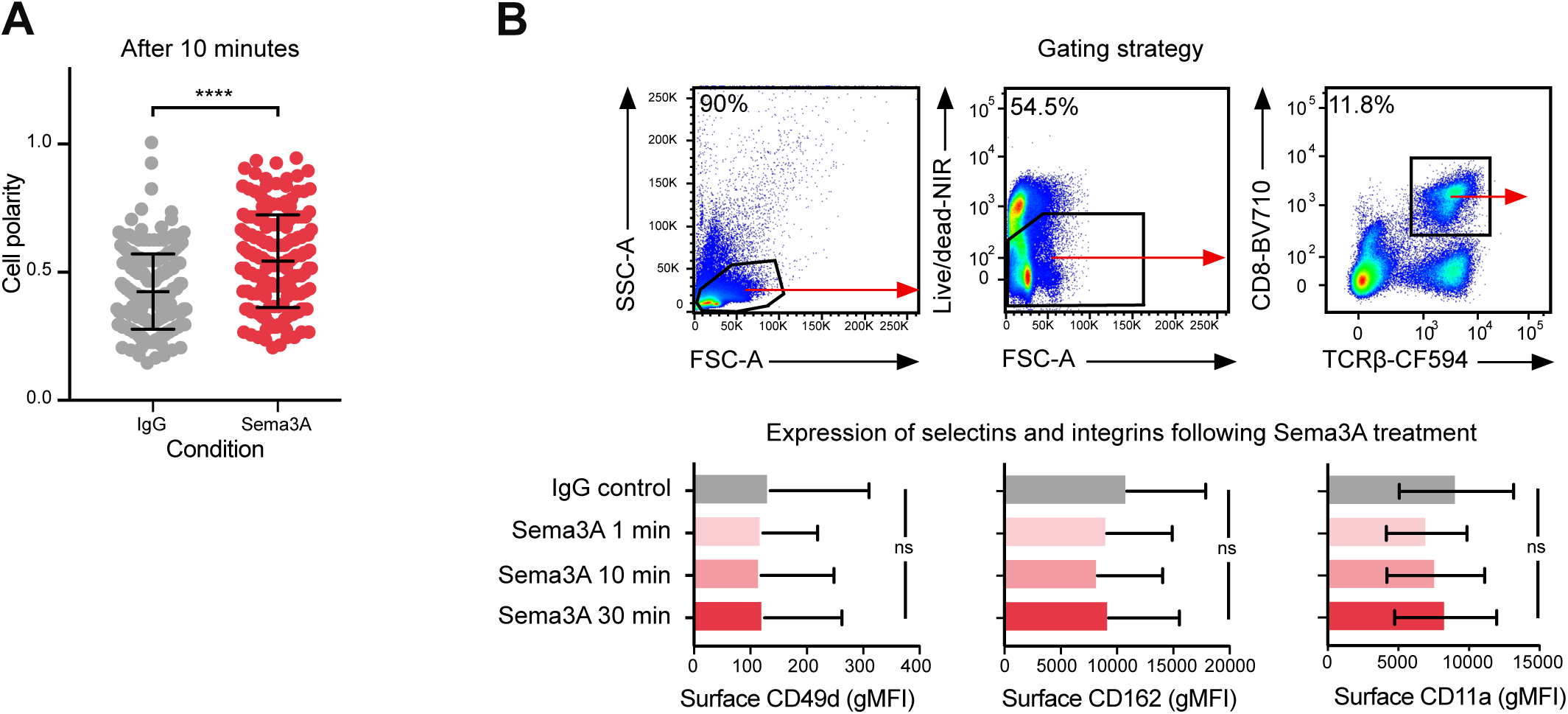
**A.** Relative frequency of cell polarity of 48 hour stimulated OT-I T cells treated with IgG or Sema3A_S-P_. A polarity of 1 indicates a shape of a perfect circle, 0 a rectangular shape. Experiment repeated three times. **** = P < 0.0001 by Student’s t-test. **B.** Gating strategy for analyzing 48 hour stimulated OT-I splenocytes treated with Sema3A_S-P_ (top). Bar graphs of gMFI of CD49d, CD162 and CD11a following Sema3A_S-P_ treatment at indicated times. ns = not significant, by Kruskal-Wallis test.

**Supplementary Figure 3 (relates to Figure 3-4).**
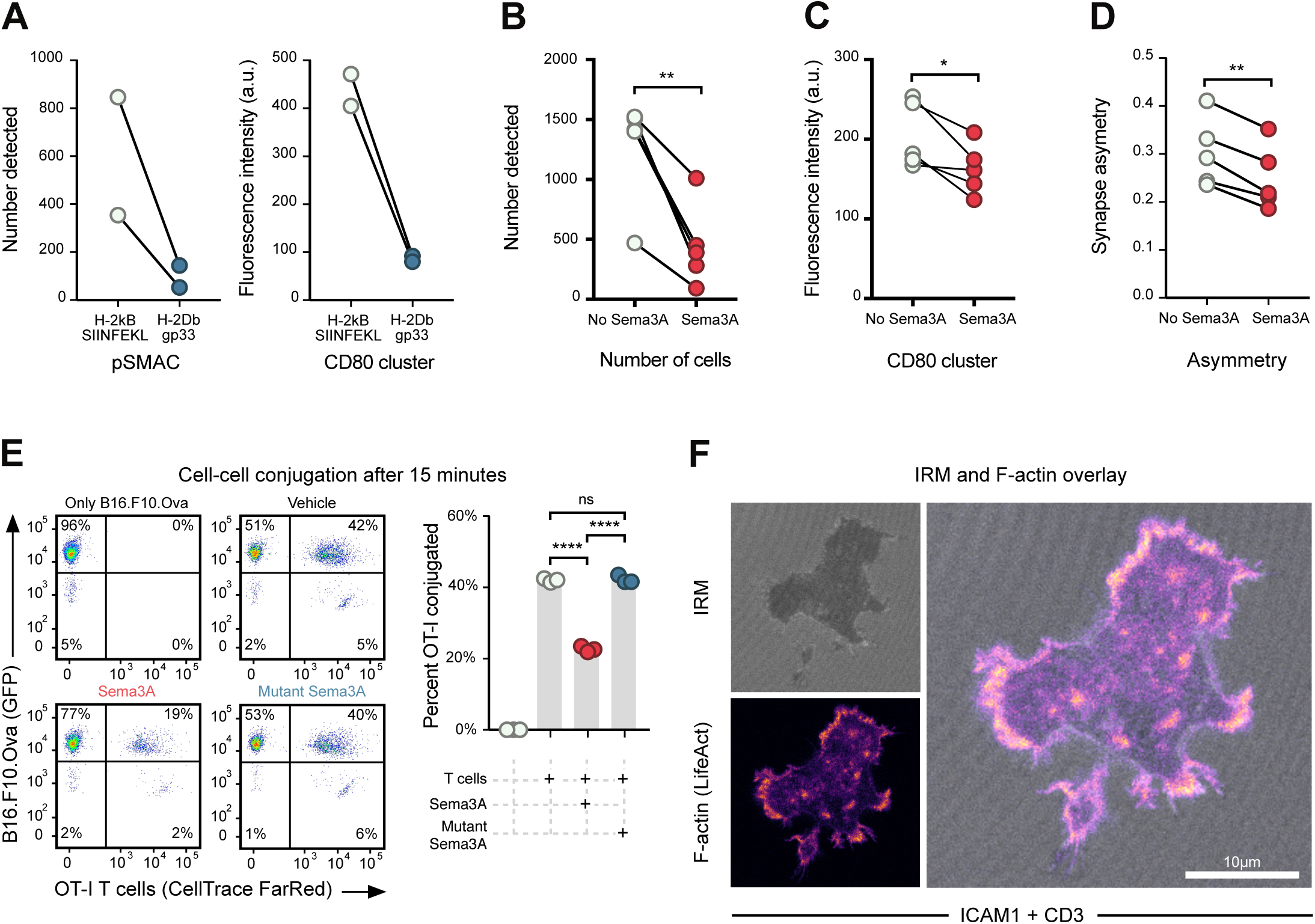
**A.** Quantification of pSMAC (left) and CD80 clustering (right) in immunological synapses of 48 hour stimulated OT-I T cells when presented with a relevant (H-2Kb-SIINFEKL) or irrelevant (H-2D-gp33) MHC monomer loaded onto the bilayer. **B.** Quantification of 48 hour stimulated OT-I T cells detected in high-throughput assay with or without Sema3A_S-P-I_ pre-treatment. ** = P < 0.01, by paired t-test. **C.** Fluorescence intensity of CD80 signal introduced by 48 hour stimulated OT-I T cells in high-throughput assay with or without Sema3A_S-P-I_ pre-treatment. * = P < 0.05, by paired t-test. **D.** Analysis of radial symmetry of synapses in OT-I T cells in high-throughput assay with or without Sema3A_S-P-I_ pre-treatment. Asymmetry of the synapse is quantified as the distance between the centroids of the CD80 cluster and that of the pSMAC relative to the diameter of the cell. ** = P < 0.01, by paired t-test. **E.** Gating strategy and representative images showing number of B16.F10.Ova cells and T cells as either singlets or doublets under four different conditions: cancer cells alone, with normal media, media with Sema3A_S-P_ or with mutant Sema3A (left). Quantification of 3 biological replicates, showing approximately 50% reduction in cell-cell conjugation when Sema3A is present (right). Data representative of three independent experiments. **** = P < 0.0001, ns = not significant, by two-way ANOVA. Gray bars indicate mean. **F.** Images from live-cell imaging of OT-I × LifeAct T cells showing concordance between IRM shadow and F-actin signal. Scalebar 10 μm. Abbreviations: a.u., arbitrary unit.

**Supplementary Figure 4 (relates to Figure 5).**
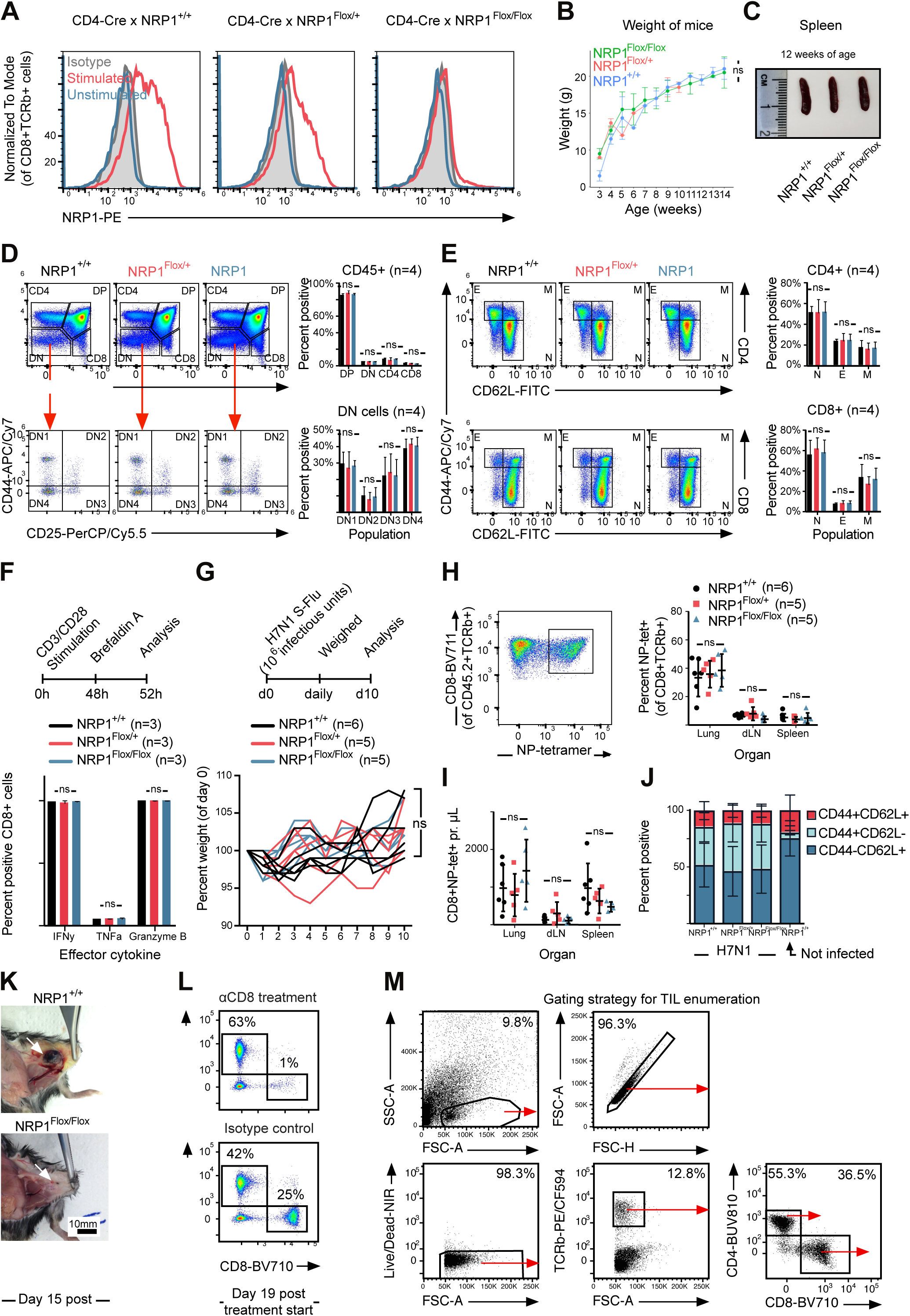
**A.** Flow cytometric analysis of naïve or CD3/CD28 stimulated splenocytes from either Nrp1^+/+^, Nrp1^Flox/+^ or Nrp1^Flox/Flox^ mice. Cells are gated on CD8 and TCRβ. Experiment performed three times. **B.** The weight of female littermates (n=17) were followed for 12 weeks and revealed no difference between genotypes. Data indicate mean ± SD. ns = not significant by two-way ANOVA. **C.** The size of spleens from female littermate mice of different genotypes at 12 weeks of age. **D.** Representative plots showing the distribution of double negative, double positive, CD4 and CD8 positive thymocytes (upper panel, left) and DN1, DN2, DN3 and DN4 populations (lower panel, left) in Nrp1^+/+^, Nrp1^Flox/+^ or Nrp1^Flox/Flox^ mice (n=4 per genotype) as analyzed by flow cytometry. Cells are gated on CD45.2. Quantification of cell populations in different genotypes (upper and lower histograms, right). Experiment performed once. ns = not significant by two-way ANOVA. **E.** Representative plots showing T cell effector populations in splenic CD4+ (top) and CD8+ (bottom) T cells. Cells are gated on CD45.2 and TCRβ (n=4 per genotype). Data combined from two independent experiments. ns = not significant by two-way ANOVA. **F.** Cytokine production following ex vivo stimulation by CD3/CD28. Experimental design (upper panel). Quantification of IFNγ, TNFα and Granzyme B by intracellular staining (lower panel). Cells are gated on TCRβ and CD8 (n=3 mice per genotype). Experiment repeated twice. ns = not significant by two-way ANOVA. **G.** Weight of mice following H7N1 S-Flu infection. Experimental design (upper panel). Weight of mice, normalized to day 0 of individual mouse weight (lower panel). Experiment performed once (n=5-6 mice per genotype). ns = not significant by two-way ANOVA. **H.** Analysis of H7N1 S-Flu-specific T cells 10 days post-infection. Example H-2D^B^-NP tetramer staining in lung of infected mouse (left figure). Quantification of H-2D^B^-NP tetramer positive CD8+ T cells in lung, dLN and spleen (right figure). Cells are gated on CD45.2, TCRβ and CD8 (n=5-6 per genotype). Experiment performed once. ns = not significant by two-way ANOVA. **I.** Quantification of total number of infiltrating H-2D^B^-NP tetramer positive CD8+ in lung, dLN and spleen 10 days post-infection. Experiment performed once. ns = not significant by two-way ANOVA. **J.** Analysis of effector subpopulations in lung 10 days post-infection in different genotypes of mice (n=5-6 mice per genotype). **K.** Representative image of B16.F10 tumors 15 days post-injection in Nrp1^+/+^ (upper image) and Nrp1^Flox/Flox^ (lower image) mice. Arrows indicates tumors. Scalebar 10 mm. **L.** Representative flow cytometric analysis of peripheral blood in mice treated with either aCD8 antibodies (upper scatterplot) or isotype control (lower scatterplot). **M.** Gating strategy used for flow cytometric analysis of TIL enumeration in mice. Abbreviations: dLN, draining lymph node. DN, double negative. E, effector T cells. N, naïve T cells. M, memory T cells. TIL, tumor-infiltrating leukocytes.

**Supplementary Figure 5 (relates to Figure 5).**
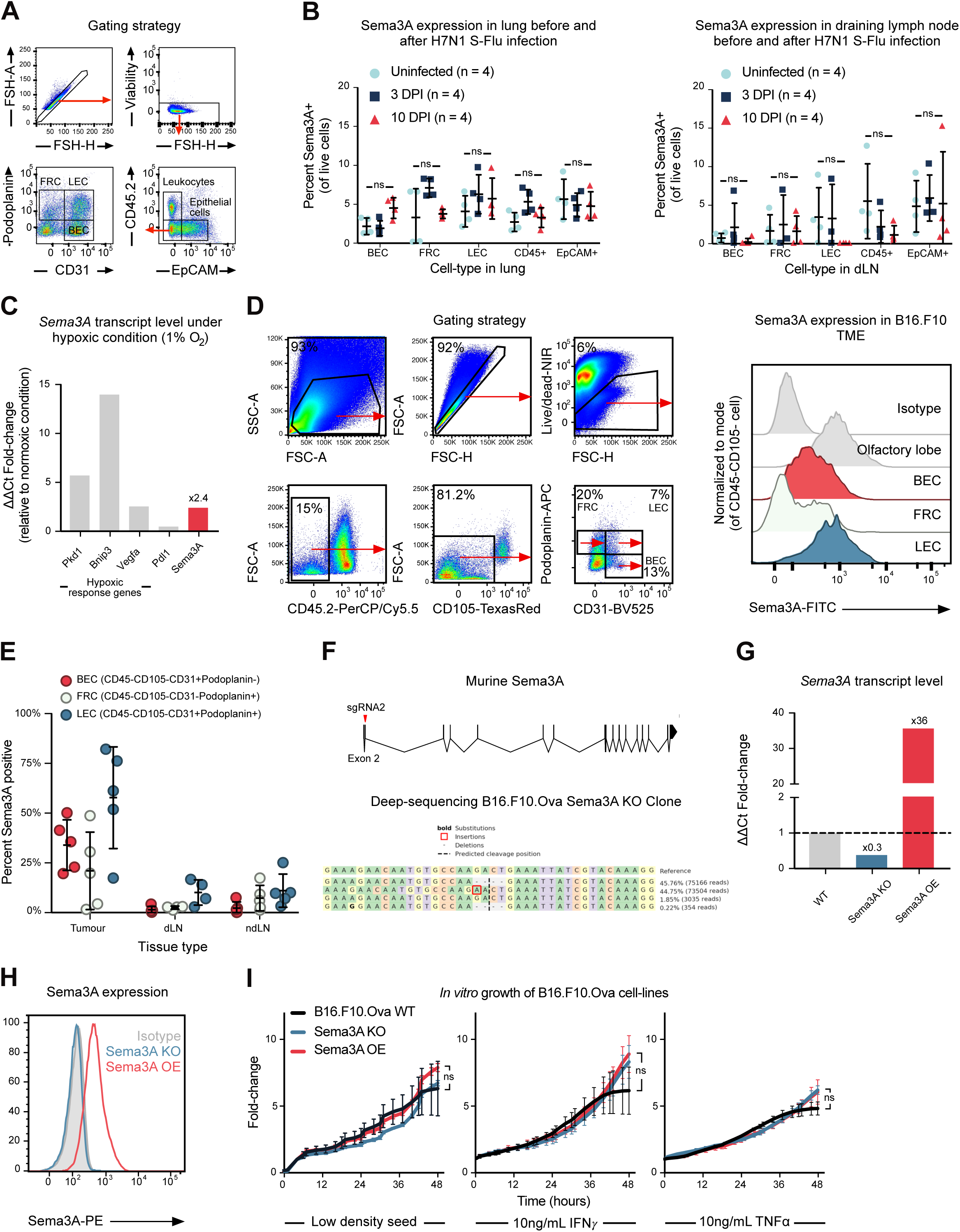
**A.** Gating strategy for analyzing Sema3A expression among leukocytes, epithelial and endothelial cells in lung and dLN. **B.** Quantification of Sema3A positive cells in different cell populations in uninfected (n=4), and infected mice, at 3 days DPI (n=4) and 10 DPI (n=4) in lung (left panel) or dLN (right panel). Nrp1^Flox/Flox^ mice used. Data combined from two independent experiments. ns = not significant by two-way ANOVA. **C.** Quantification of Pkd1, Bnip3, Vegfa, Pdl1 and Sema3A mRNA level following 24 hour culture in 1% O2 chamber. **D.** Gating strategy (left) and representative histograms (right) analyzing Sema3A positive cell populations in B16.F10 tumors 11 days post-injection. Olfactory lobe is used as a positive control for Sema3A expression. **E.** Quantification of Sema3A positive cells in same experiment as in (D) in tumor, dLN and ndLN. **F.** Genomic organization of murine Sema3a gene, indicating where CRISPR guide RNA targets with red arrow (upper figure). MiSEQ sequence results for chosen Sema3A KO clone showing 4 base deletion in two alleles (46% of all reads), a frameshift in one allele (45% of all reads) and WT reads in 2% of all reads. **G.** RT-qPCR analysis show down- and up-regulation of Sema3A in Sema3A KO and OE cell lines, respectively. Normalized to Hprt. Experiment performed once. **H.** Intracellular staining shows no detectable expression of Sema3A in Sema3A KO cells, and expression in Sema3A OE cells, as expected. Experiment performed once, at low seeding density. **I.** Growth of WT, Sema3A KO and Sema3A OE B16.F10.Ova cell lines in normal, IFNγ or TNFα-rich media. Experiment performed once. Data indicate mean ± SD. ns = not significant by two-way ANOVA. Abbreviations: BEC, blood endothelial cells. dLN, draining lymph node. DPI, days post-infection. FRC, fibroblastic reticular cells. KO, knockout. LEC, lymphatic endothelial cells. ndLN, non-draining lymph node. OE, overexpressing. TME, tumor microenvironment. WT, wild-type.

**Supplementary Figure 6 (relates to Figure 6).**
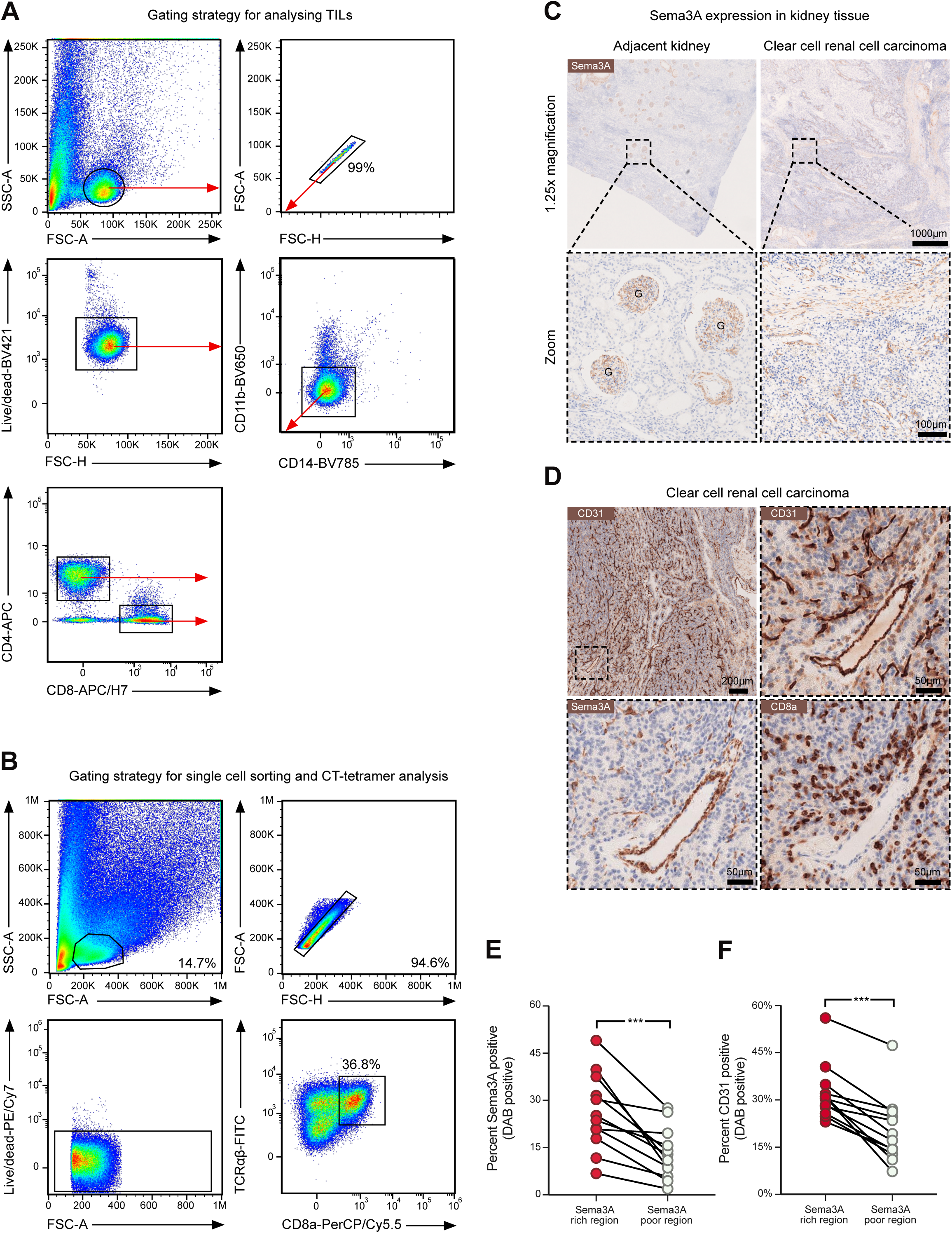
**A.** Gating strategy for analyzing TILs from ccRCC patients. **B.** Gating strategy for single cell sorting and CT-tetramer analysis. **C.** Sema3A expression in tumor-adjacent and tumor tissue of ccRCC patient. G indicates kidney glomeruli. Scalebars indicate 1000 μm (upper row) and 100 μm (lower row). **D.** Serial sections from ccRCC tumor stained for CD31, Sema3A and CD8a. Dashed box in upper left image indicates the region depicted at higher magnification in the three other images. Scalebar indicates 200 μm (upper left) and 50 μm (other images). **E.** Expression of CD31 in Sema3A rich and poor regions. **** = P < 0.0001 by paired t-test. **F.** Expression of Sema3A in selected Sema3A rich and poor regions. **** = P < 0.0001 by paired t-test. Abbreviations: ccRCC, clear cell renal cell carcinoma. DAB, 3,3’-Diaminobenzidine.

